# Glutamylated vimentin proteoform identified by a synthetic binder links to epithelial-mesenchymal plasticity

**DOI:** 10.64898/2026.05.12.724742

**Authors:** Rayees A. Ganie, Anwesha Biswas, Sarika Sasi, Lakshmi Prasanna Devarakonda, Ishan Kale, Sanjana Mullick, Gangotri Siddappa, Nirpendra Singh, Sudarshan Gadadhar, Minhajuddin Sirajuddin

**Affiliations:** Institute for Stem Cell Science and Regenerative Medicine (iBRIC-inStem), GKVK Campus, Bengaluru – 560065, India; Manipal Academy of Higher Education, Manipal, Karnataka, 576104, India; Regional Centre for Biotechnology (RCB), NCR Biotech Science Cluster, 3rd Milestone, Faridabad-Gurugram Expressway, Faridabad, 121001, Haryana (NCR Delhi), India

## Abstract

Cytoskeletal protein expression and filament dynamics change significantly during cell state transitions. However, post-translational modifications of cytoskeletal proteins during these transitions have rarely been described. Here, using a synthetic glutamylation-binder (SB2B49) selected against a bi-glutamylated peptide epitope, we identify a distinct pool of glutamylated vimentin filaments. We demonstrate that vimentin glutamylation is enzymatically added by tubulin tyrosine ligase-like (TTLLs) and removed by cytosolic carboxypeptidases (CCPs). Mass spectrometry and mutagenesis reveal that glutamylation occurs on specific vimentin residues. We find that glutamylated vimentin levels are dynamically modulated during epithelial-mesenchymal plasticity. During collective migration in scratch-wound assays, glutamylated vimentin filaments are transiently depleted at the wound edge, a process controlled by canonical glutamylation writer-erasers. Our findings reveal a new layer of vimentin regulation via glutamylation and establish the glutamylation-binder as a valuable tool for exploring the diversity of vimentin proteoforms and glutamylation modifications in both physiological and pathological contexts.

## Introduction

Cytoskeletal polymers such as microtubules, actin filaments, and intermediate filaments serve functions beyond merely providing structural support. They are dynamic filaments that influence cell mechanics, motility, stability, and signaling^1,2^. A growing body of evidence shows that these filaments can be divided into molecularly distinct subpopulations that support specialized functions in different spatial and temporal contexts. For microtubules, this diversification is often described as tubulin code^3^, shaped by tubulin isotypes and post-translational modifications (PTMs) such as acetylation^4^, detyrosination/tyrosination cycle^5,6^, and glutamylation^7^. These features tune microtubule interactions with motors^8,9^ and MAPs^10^ and are linked to changes in cell organization, division^11^, and transport^12^. Similar principles are emerging for actin, where actin isoforms and PTMs extend functional states beyond a single generic filament^13,14^. In comparison, although intermediate filaments (IFs) are key determinants of cellular and tissue mechanics, how PTMs differentiate IF networks into functionally distinct proteoforms remains poorly understood^15–17^.

Vimentin is a major IF protein that supports cell shape, mechanical resilience, and cytoskeletal integration. Its expression increases during epithelial-mesenchymal transition (EMT) and is widely used as a marker of mesenchymal, invasive, and metastatic cell states^16,18–20^. EMT itself is a reversible cell-state program in which epithelial cells lose polarity and junctional organization while adopting mesenchymal behaviors such as increased motility and invasiveness. EMT (and partial/hybrid EMT states) is important in development and wound repair, can contribute to fibrosis, and is frequently co-opted in cancer progression and metastasis^21–23^. Canonical EMT-inducing cues, including TGF-β, EGF, FGF, and IGF signaling, converge on transcriptional regulators such as SNAIL, TWIST, and ZEB family factors to rewire gene expression and cytoskeletal organization^16,18,21,24,25^. Consistent with vimentin’s central role in these transitions, vimentin is extensively modified by PTMs, most prominently multisite phosphorylation, but additional modifications are reported across contexts. Yet, how specifically PTM-defined vimentin proteoforms regulate filament stability, remodeling, and EMT phenotypes remains an open question^15,16,26^.

Among cytoskeletal PTMs, glutamylation is especially well characterized in the microtubule field because the glutamylases (writers) and deglutamylases (erasers) were identified early^3,27–30^, and robust antibodies enabled visualization and biochemical analysis of glutamylated tubulin^31,32^. The writer enzymes (tubulin tyrosine ligase–like proteins; TTLLs) can catalyze initiation and elongation of glutamate side chains, while the erasers (cytoplasmic carboxypeptidases; CCPs) family members remove these chains, together shaping glutamylation patterning and signaling capacity on tubulin^29,30^. At the same time, glutamylation is *not* restricted to tubulin, and growing evidence suggests that TTLL/CCP biology cannot always be explained by microtubule regulation alone^3,33^. A practical bottleneck has been the limited availability of probes that can specifically distinguish modified from unmodified proteoforms in cells, especially in formats compatible with live-cell reporting and with direct readouts of enzymatic activity rather than endpoint staining.

Synthetic binders provide a flexible route to the engineering of PTM-selective affinity reagents and intracellular sensors. Here, we developed a glutamylation binder by screening a yeast-display SSO7D library against a common glutamylation epitope. This approach yielded SB2B49, a synthetic binder that robustly reports glutamylated microtubules across cell types, highlighting centrosomes/centrioles, spindle poles, and the midbody, as well as glutamylation-rich structures such as basal bodies and axonemes. Unexpectedly, SB2B49 also labels filamentous structures that co-localize with the glutamylation antibody GT335. These were then identified as vimentin IFs by multiple orthogonal validations. Notably, SB2B49-marked glutamylated vimentin correlates with epithelial-like gene profiles and diminishes during EMT. Together, these results establish a general platform for discovering glutamylation-defined proteoforms in cells and reveal vimentin glutamylation as a regulated cytoskeletal feature.

## Results

### Screening for glutamylation binders

A central challenge in studying glutamylation outside of tubulin has been the scarcity of probes that distinguish modified from unmodified proteoforms with sufficient specificity in cells. Similar to our previous approach in developing the microtubule tyrosination sensor^34^, we chose the tubulin carboxy-terminal tail peptide epitope bearing the glutamylation modification for screening against the SSO7D binder yeast display library. Biotinylated Hs_TUBB2 427-445-biE [435], herein referred to as Beta-2-BiE, represents the most common and abundant glutamylation modification^35^. A similar epitope was used to generate GT335 and Beta-monoE, the most widely used glutamylation antibodies in the field^31,32^. The biotinylated-Beta-2-BiE peptide was used to enrich for potential SSO7D glutamylation binders via MACS- and FACS-based screening (Methods) (Fig. 1a, 1b, and Extended Data Fig. 1a).

**Figure 1:**
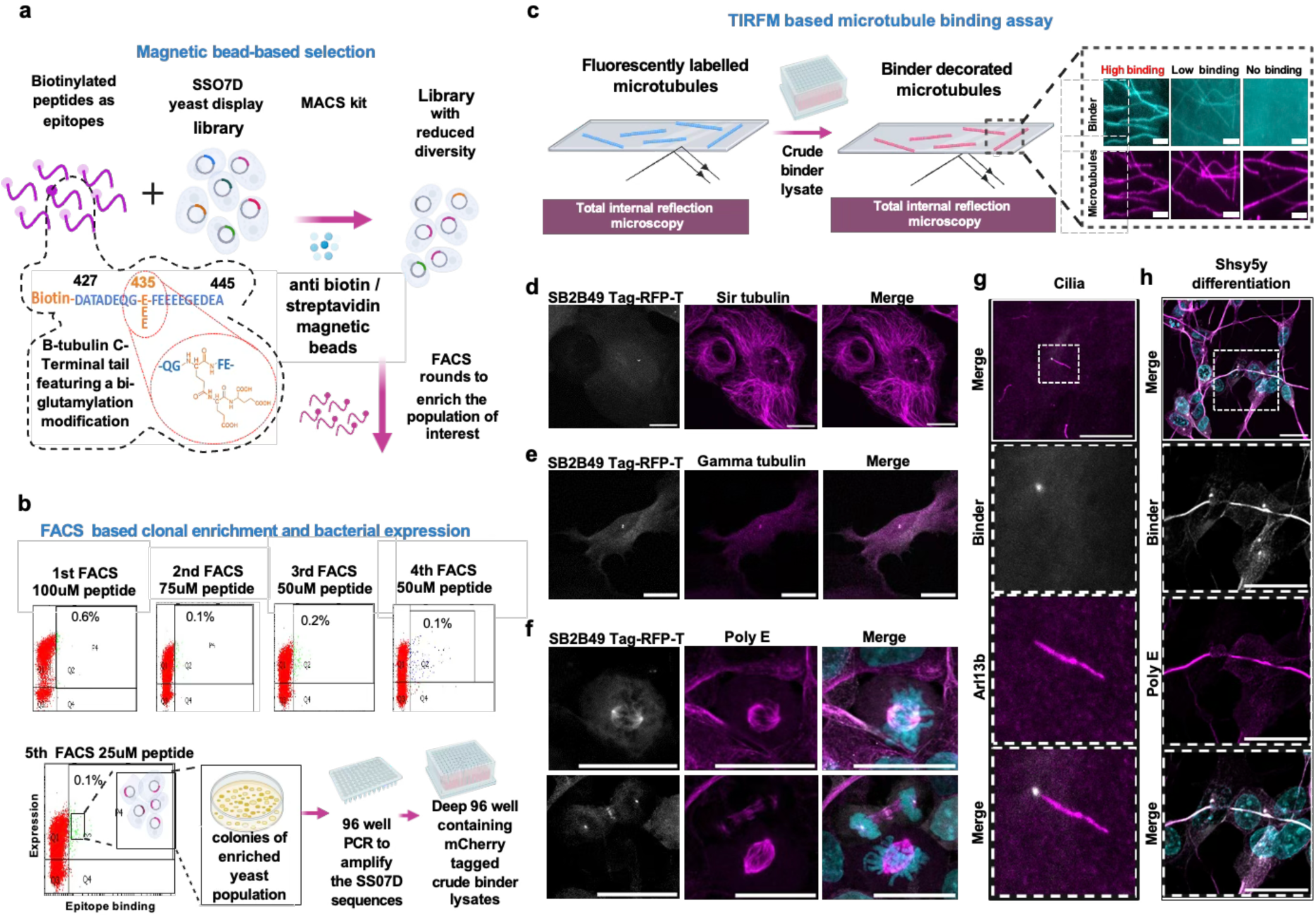
A platform for screening, selection, and characterization of a glutamylation biosensor. **a-c.** Schematic of the binder discovery workflow. A biotinylated beta-tubulin carboxy-terminal tail peptide bearing a defined bi-glutamylated epitope (B2-BiE) was used to screen a yeast-display SSO7D binder library. Magnetic bead-based enrichment (MACS) was followed by iterative FACS selections using decreasing B2-BiE concentrations, with enrichment monitored as the fraction of double-positive yeast. After the fifth FACS round, individual clones were sequenced, subcloned into a bacterial expression vector, and expressed in 96-well format. Candidate binders were screened by TIRF microscopy for binding of mCherry-tagged binders to purified brain microtubules, and high-binding candidates were advanced to cell-based validation. **d.** Representative confocal images of U2OS live cells stably expressing SB2B49–TagRFP-T (grayscale) with SiR-tubulin (magenta). **e.** Immunostained confocal images of RPE1 cells stably expressing SB2B49–TagRFP-T (grayscale) with Gamma-tubulin antibody (magenta). **f.** Confocal images of stable SH-SY5Y cells expressing SB2B49–TagRFP-T (grayscale), immunostained with PolyE antibody (magenta), and imaged in different cell-cycle stages. DNA was labeled with Hoechst (cyan). **g.** Ciliary microtubule glutamylation in RPE1 cells. Representative images of ciliated RPE1 cells stably expressing SB2B49–TagRFP-T (grayscale), fixed and stained for Arl13b (magenta). **h.** Axonal microtubule labeling during neuronal differentiation. Representative images of differentiated SHSY5Y cells stably expressing SB2B49–TagRFP-T (grayscale) and immunostained with PolyE antibody (magenta). The white-dotted box represents the zoomed-in ROI region as indicated. Scale bars = 20 μm (unless otherwise indicated).

Post-screening, 44 individual yeast colonies were cloned into a bacterial expression vector and sequenced to analyze enrichment of binder clones (Methods) (Fig. 1a, 1b and Extended Data Fig. 1b). Sequence alignment revealed 32 unique SSO7D clones, which were then expressed in bacteria as mCherry-fused proteins and tested in a microtubule binding assay using TIRF microscopy (Methods) (Fig. 1c, Extended Data Fig. 1b and 2a). Based on the TIRF images, we classified the SSO7D clones into microtubule-strong, low-, and non-binders (Fig. 1c and Extended Data Fig. 2a). Eight strong microtubule-binding SSO7D clones were evaluated for their specificity towards unmodified and glutamylated microtubules^36^. Microtubules were polymerized individually from HEK tubulin and goat brain tubulin, mixed with Cy5-labelled tubulin, representing the unlabeled unmodified and Cy5-glutamylated brain microtubules within the same flow chamber (Methods) (Extended Data Fig. 2b). mCherry-SSO7D binders were flowed into the chambers containing the dual set of microtubules and imaged using TIRF microscopy (Methods). The TIRF images show that 6 out of 8 strong microtubule-binding SSO7D clones specifically bind to glutamylated brain microtubules (Extended Data Fig. 2c).

**Figure 2.**
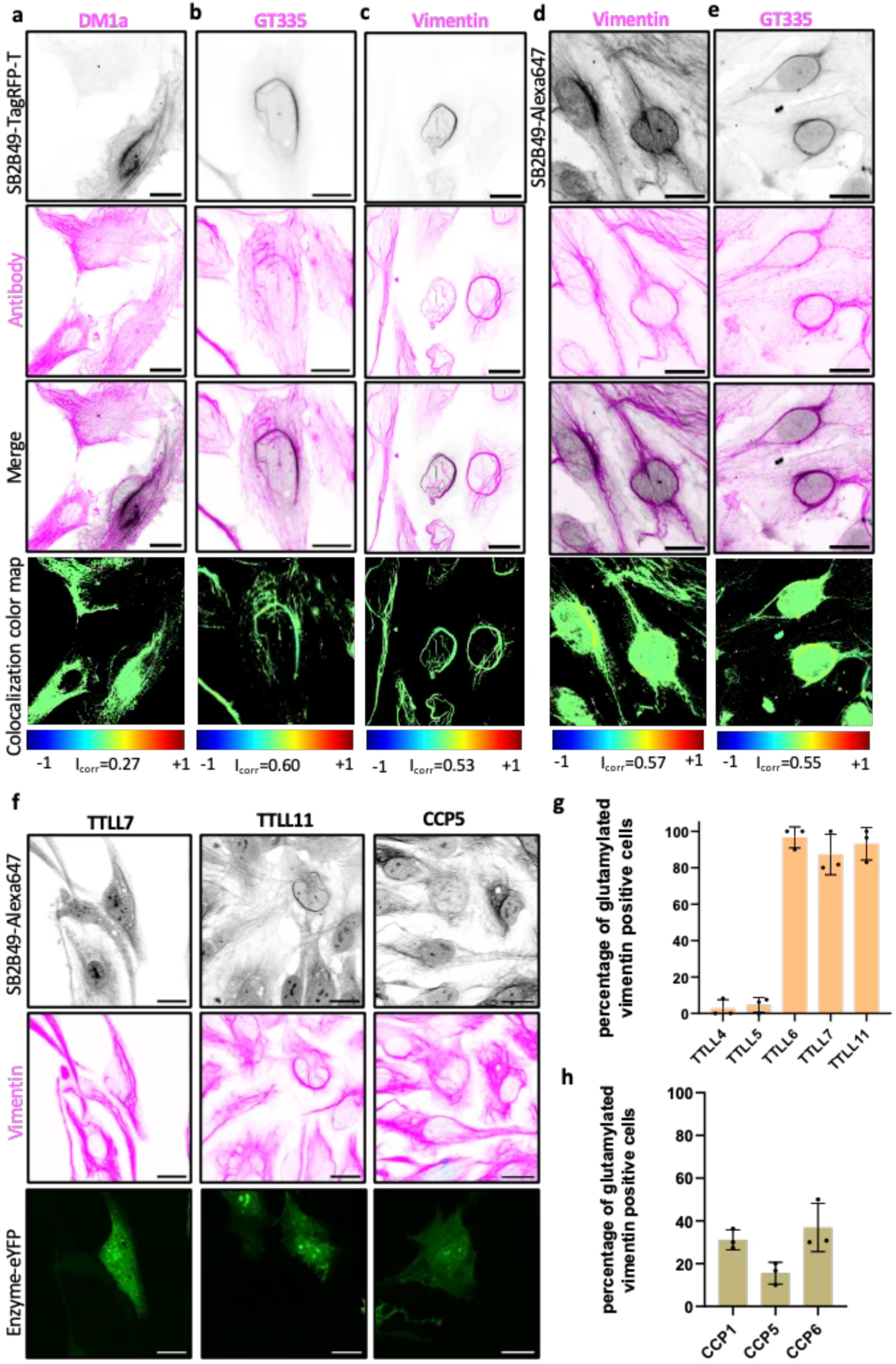
Discovery of a unique subset of vimentin by the glutamylation sensor. **a – c** Representative images of RPE1 cells stably expressing Tag-RFP-T fused to SB2B49 (grayscale), microtubules stained with DM1a or vimentin antibody or GT335 (magenta) as indicated by Pearson correlation coefficient data. The presence of a unique perinuclear structure in a subset of cells is marked by the yellow arrow and the centrosome with the red arrow. **d & e**. Confocal images of fixed and stained RPE1 cells with recombinant SB2B49 binder labelled with CF649 (grayscale) and GT335 or vimentin (magenta) antibody as indicated. **f.** Representative confocal images of stable RPE1 cell line co-expressing hTTLL7-, hTTLL11, CCP5-eYFP (green), and Tag-RFP-T fused to S2B49 binder (grayscale), fixed and stained with GT335 (magenta). g Scale bars = 20 μm **g & h.** Quantification of the percentage of cells expressing hTTLL4-, hTTLL5-, hTTLL6-, hTTLL7-, hTTLL11, CCP1-, CCP5-, CCP6-eYFP that have glutamylated vimentin signal from SB2B49 binder labelled with CF649. Error bars represent the mean with SD.

### Glutamylation binders bind to *bona fide* glutamylation structures in cells

The six SSO7D binders were then fused to TagRFP-T and transiently expressed in U2OS cells, and the cells were imaged with SiR-tubulin (Methods). Most of the SSO7D binders showed diffusive cytoplasmic signal, indicating no binding to microtubules or any subcellular structures. However, four SSO7D binders showed a strong binding signal to subcellular structures. SB2B34 and SB2B36 were localized to the nucleus and showed bright patches inside the nucleus. SB2B49 and SB2B60 exhibited a strong speckle-like signal near MTOC and a subset of filamentous structures adjacent to the nuclei across different cell lines (Fig. 1d, Extended Data Fig. 3a and 3b). Since cells generally lack glutamylation during interphase, except at centrosomes/centrioles, we examined whether SB2B49 localizes to them. Immunostaining experiments with the anti-gamma-tubulin antibody and SB2B49-TagRFP-T show fluorescence colocalization, indicating that SB2B49 indeed marks the centrosome/centrioles in U2OS and RPE1 cells (Fig. 1e and Extended Data Fig. 3c).

**Figure 3.**
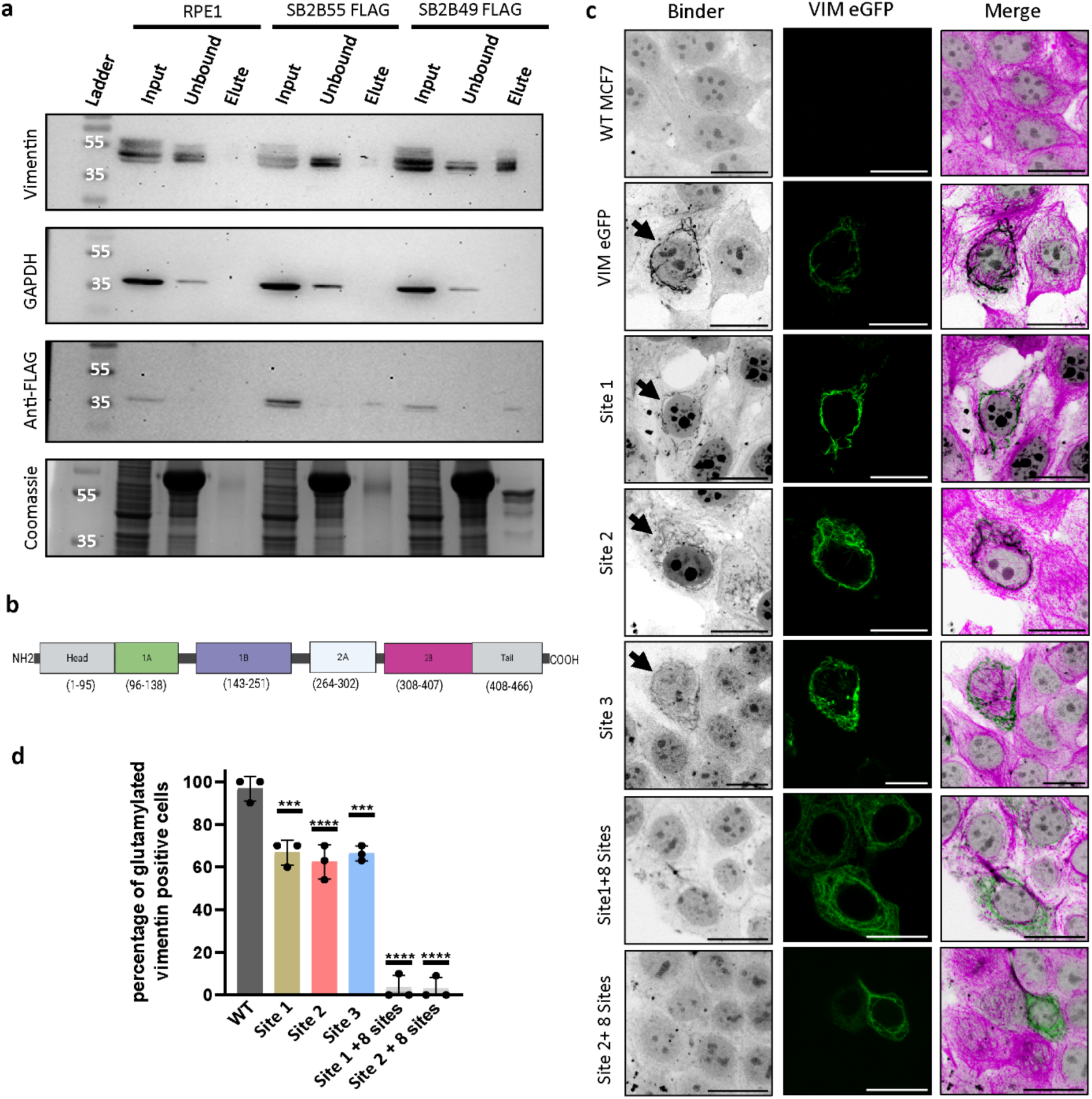
Mapping and functional testing of vimentin glutamylation sites. **a.** Cell lysates from stably expressing SB2B55 (Random binder) and SB2B49 (Glu-binder) tagged with FLAG peptide, and wild-type RPE1 cells were fractionated using anti-FLAG beads (Methods). The input, unbound, and eluted fractions were probed using anti-vimentin, GAPDH, and FLAG antibodies using a Western blot. The same fractions were simultaneously analyzed by SDS-PAGE with Coomassie stain. Only the SB2B49 (Glu-binder) elute fraction yields a positive signal on Western blot probing with anti-vimentin. **b.** Domain organization of vimentin . **c.** Confocal images of MCF7 cells expressing WT and Glu-mutant Vimentin-eGFP (green), Glu-binder (grayscale), as indicated. Scale bars = 20 μm **d.** Quantification of the percentage of MCF7 cells expressing vimentin wildtype and Glu-mutants, as indicated, has a glutamylated vimentin signal from SB2B49 binder labeled with CF649. Sites 1, 2 , and 3 abolish Glu-binder binding compared to WT. Statistical significance was determined with Ordinary one-way ANOVA. Each point represents an independent biological replicate. Error bars represent mean with SD, ****= P value <0.0001, ***= P value <0.0002.

We next followed the SB2B49 TagRFP-T fluorescence signal across different stages of the cell cycle and found that it marks *bona fide* glutamylation structures during mitosis, such as the centrosomes/centrioles during interphase, spindle poles from Prophase to Telophase, and mid-body microtubules during cytokinesis (Fig. 1f and Extended Data Fig. 3d). The SB2B49 TagRFP-T also marks the basal body of cilia (Fig. 1g and Extended Data Fig. 3e). However, the recombinant SB2B49 binder marks only the axonemal structures of mouse sperm (Extended Data Fig. 3f). This could be due to the SB2B49 TagRFP-T fusion protein being unable to transit into the cilia in live cells, or the mouse sperm axonemal microtubules being biochemically distinct, which requires further investigation. Another prominent glutamylation modification site is the axons of neurons. Immunostaining experiments with SB2B49 TagRFP-T and glutamylation antibodies show that only a subset of axons is labeled by the SB2B49 TagRFP-T signal. This indicates that the SB2B49 can be used to track axonal microtubules (Fig.1h and Extended Data Fig. 3g) In summary, SB2B49 marks *bona fide* microtubule glutamylation structures, and henceforth, we refer to it as a glutamylation sensor.

### Identification of vimentin glutamylation modification

We then assessed whether SB2B49 and SB2B60 bind to glutamylated microtubules following ectopic expression of glutamylases. Overexpression of glutamylases (TTLL4, 5, and 7), the SB2B49 and SB2B60, failed to mark any cytoplasmic microtubules (Extended Data Fig. 4a). This could largely be because overexpression of glutamylases can modify multiple glutamate residues in the CTT of tubulin^37^, and SB2B49 and SB2B60 could be sensitive to such molecular alterations at the tubulin carboxy-terminal tail.

The glutamylation sensor marks the centrosome/centriole in interphase cells (Fig. 1d, 1e, and Extended Data Fig. 3c). We also observed filamentous structures marked by the fluorescence of the glutamylation sensor in a few cells (Fig. 2a, and Extended Data Fig. 3a and 4b). These structures did not colocalize with known microtubule-specific antibodies or their modifications, except for GT335 (Fig. 2a, 2b, and Extended Data Fig. 4c).

**Figure 4.**
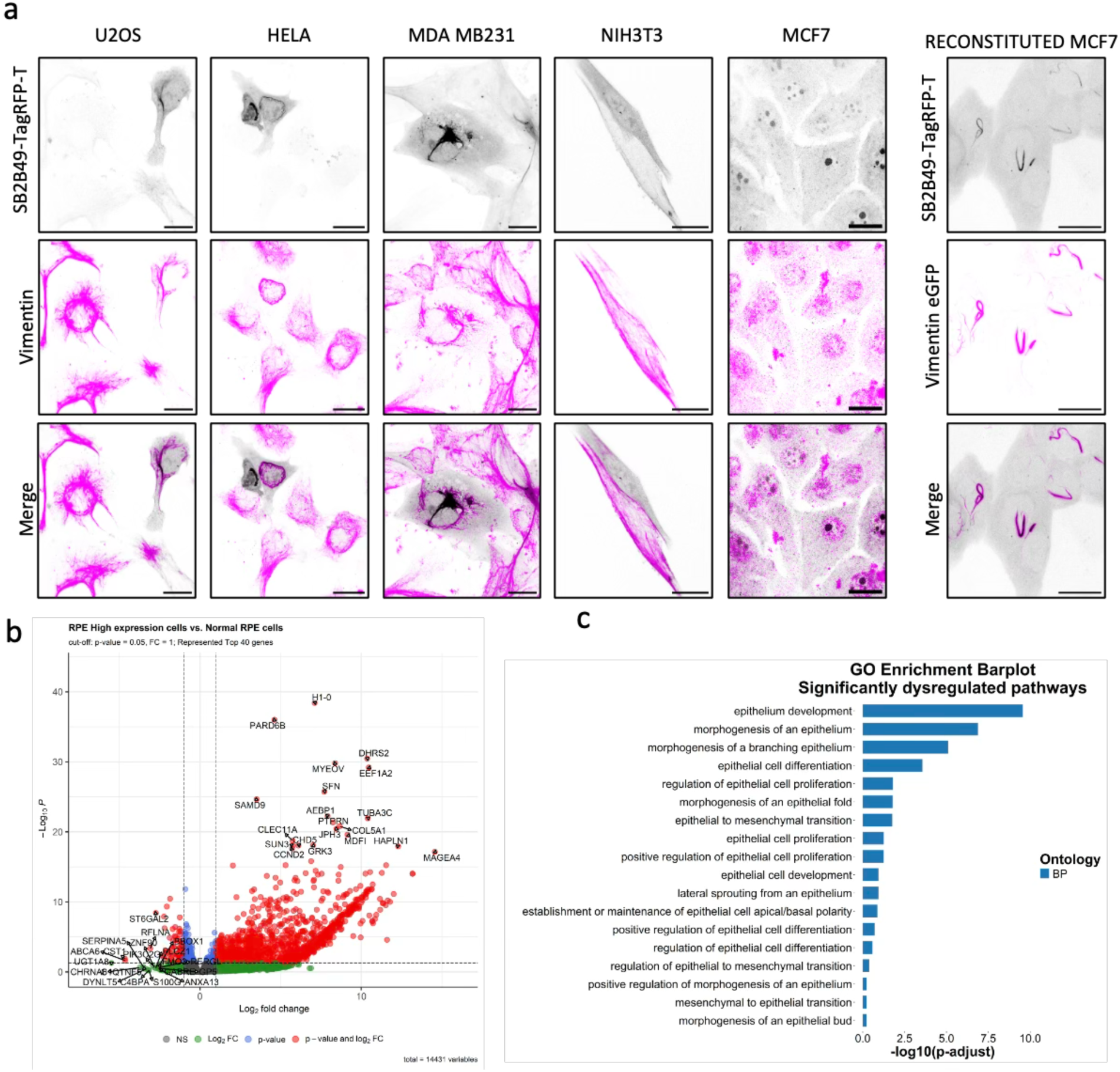
Glutamylated vimentin is associated with the epithelial state of the cells. **a.** Different cell lines expressing SB2B49 binder fused to Tag-RFP-T (grayscale) were fixed and stained with anti-vimentin antibody (magenta). Notably, the mesenchymal fibroblast cell line NIH3T3 does not show the presence of the modified vimentin structure. As expected, MCF7 cells, which are devoid of any vimentin, also do not show the presence of the vimentin structure. Representative images of double stable MCF7 cells co-expressing SB2B49 binder fused to Tag-RFP-T (grayscale) and eGFP tagged vimentin (magenta), showing the labelling of a subset of the overexpressed vimentin. Scale bars = 20 μm **b.** volcano plot showing differences in epithelial and mesenchymal gene expression between RPE1 high and control. **c.** RNA-seq data showing gene ontology and expression differences for a subset of genes associated with epithelial and mesenchymal states in RPE1 high cells compared to controls.

To identify the molecular composition of the filamentous structures, we counterstained the TagRFP-T fused glutamylation sensor against vimentin, keratin, and the desmin cytoskeleton (Fig. 2c and Extended Data Fig. 4d) (Methods). Among the cytoskeletal filaments staining, only vimentin colocalized with the glutamylation sensor and the GT335, but not with DM1A (Fig. 2a-2c). To rule out the possibility that the filamentous structures are artifacts of the glutamylation sensor labeling, we stained the cells with recombinant SB2B49 labeled with CF640R, hereon referred to as recombinant glutamylation binder (Methods). The staining confirmed that the filamentous structures are inherent to the cells, regardless of ectopic expression of the fluorescent-tagged glutamylation sensor (Fig. 2d and 2e). We then introduced a series of known glutamylases (TTLL5, 6, 7, and 11) and checked whether vimentin glutamylation was affected (Methods). Fluorescent images of the recombinant glutamylation binder show that TTLL6, 7, and 11 glutamylases increase glutamylated vimentin levels compared to other glutamylases (Fig. 2f, 2g, and Extended Data Fig. 5a). On the other hand, expression of deglutamylases CCP1, 5, and 6 abolishes the recombinant glutamylation binder fluorescence on the vimentin filaments (Fig. 2f, 2h, and Extended Data Fig. 5a).

These experiments strongly suggest the presence of glutamylated vimentin proteoform in cells, as detected by the glutamylation sensor, binder, and the GT335 antibody. Additionally, cells use canonical glutamylation writers and erasers to modify vimentin filaments.

### Validation and characterization of glutamylated vimentin modification

To biochemically confirm the presence of glutamylated vimentin in cells, we performed glutamylation sensor pull-down assays (Methods). Coomassie staining and vimentin Western blot analysis of RPE1 cells with FLAG-tagged glutamylation sensor show a clear vimentin enrichment compared to the control set (Fig. 3a and Extended Data Fig. 6a-c).

The enriched pull-down fraction was subjected to proteomic analysis, which further confirmed the presence of vimentin and provided clues about potential glutamylation sites (Methods) (Fig. 3a, Extended Data Fig. 6d). The vimentin glutamylation sites identified by mass spectrometry showed at least 7 potential modification sites (Fig. 3b and 3c). We further identified 7 additional potential glutamylation sites that contain a stretch of glutamates resembling the tubulin carboxy-terminal tail (Fig. 3b). These glutamates were mutated to aspartates, preserving the acidic nature of the sites while preventing glutamylation^38^. To validate the vimentin glutamylation sites, we used MCF7 cells, which do not express vimentin^39,40^ (Fig. 3c), and reconstituted filamentous vimentin via ectopic expression of wild-type or mutant vimentin genes (Methods) (Fig. 3c). MCF7 cells expressing wild-type and vimentin mutated at the mass spectrometry identified sites (Site 1, Site 2 and Site 3) show filamentous vimentin (Fig. 3c, and 3d). However, the glutamylated vimentin signal using the recombinant glutamylation binder decreases with the aspartate mutations at Site 1, Site 2, and Site 3 (Methods) (Fig. 3c and 3d). To determine whether multiple glutamylation sites co-exist in vimentin, we used a combinatorial mutation approach. First, we mutated a stretch of glutamates across the vimentin primary sequence that were not identified in our mass spectrometry analysis (Vim-7-mut). Vim-7-mut and the Site 3 combination also showed a significant decrease in the reduction of glutamylated vimentin signal (Fig. 3d). Therefore, we subsequently mutated Vim-7-mut+Site 3 (Vim-8-mut) and at either the Site 1 (Vim-8-mut+Site 1) or Site 2(Vim-8-mut+Site 2) sites, which completely abolished the glutamylated vimentin signal using the recombinant glutamylation binder (Fig. 3c and 3d).

Although we do not currently understand the cooperativity between glutamylation sites and specificity towards TTLLs, our results unequivocally suggest the presence of vimentin glutamylation modification and potential vimentin glutamylation sites.

### Vimentin glutamylation modification is associated with the epithelial cell state

The vimentin cytoskeleton is a key marker during the epithelial-mesenchymal transition and vice versa^18^. To determine if vimentin glutamylation correlates with cell state, we first assessed the presence of glutamylated vimentin in various cell lines, including U2OS, RPE1, HeLa, MDA-MB-231, NIH/3T3, and MCF7 (Methods). Except for MCF7, all cell lines showed a subset of glutamylated vimentin, detected with the glutamylation sensor (Fig. 4a). We previously showed that MCF7 cells lack a vimentin cytoskeleton, and a subset of ectopically expressed vimentin-EGFP in MCF7 can be detected by the glutamylation sensor (Fig. 3d) (Methods). These results suggest that the glutamylated form of vimentin is conserved across various cell types, and the cells utilize existing glutamylation enzymes to modify vimentin.

Next, we investigated whether vimentin glutamylation is associated with a specific cell state through transcriptome analysis. RPE1 cells stably expressing glutamylation sensor were sorted based on the TagRFP-T fluorescence intensity (Methods). The fluorescence intensity of glutamylation sensor indicates increased levels of glutamylated vimentin (RPE_Vim-glu_) (Extended Data Fig. 7a). Additionally, we observed that RPE1 cells seeded immediately after thawing a frozen stock (RPE_fresh-seed_) do not contain any glutamylated vimentin (Extended Data Fig. 7b). However, the same batch of cells, after 2 passages, shows glutamylated vimentin (RPE_passaged_) (Extended Data Fig. 7b). RNA transcripts were extracted from RPE_normal_, RPE_Vim-glu_, RPE_fresh-seed_, and RPE_passaged_ cells, and their transcription profiles were compared using RNA sequencing (Methods).

Transcriptomic analysis of RPE_Vim-glu_ showed up-regulation of epithelial-related *(EPGN, ID2, LIN7A, MTSS1, SOX-2)* and EMT-related *(COL1A1, EMP2)* genes as compared to unsorted RPE_normal_ (Fig. 4b). The ontology enrichment analysis also indicated up-regulation of epithelial state in RPE_Vim-glu_ cells (Fig. 4c). Similar transcription profiles were observed when comparing RPE_fresh-seed_ and RPE_passaged_ (Extended Data Fig. 7c). Thus, corroborating our observation that glutamylated vimentin is associated with the epithelial state of the cell and can be a dynamic feature.

### Cell-state transitions dynamically modulate glutamylated vimentin and migration

The transcriptomic and cell-state analyses suggest that vimentin glutamylation is associated with the epithelial cell state. We then investigated whether external stimuli that induce cell-state transitions could affect glutamylated vimentin levels. TGF-beta1 is a well-known factor that induces epithelial-mesenchymal transition^24,41^. RPE1 cells stably expressing the glutamylation sensor were treated with TGF-beta1 and monitored after 5 days (Fig. 5a) (Methods). Confocal images of TGF-beta1-treated versus untreated cells show a clear decrease in the glutamylated form of vimentin (Fig. 5b, 5c, and Extended Data Fig. 8a-h).

**Figure 5.**
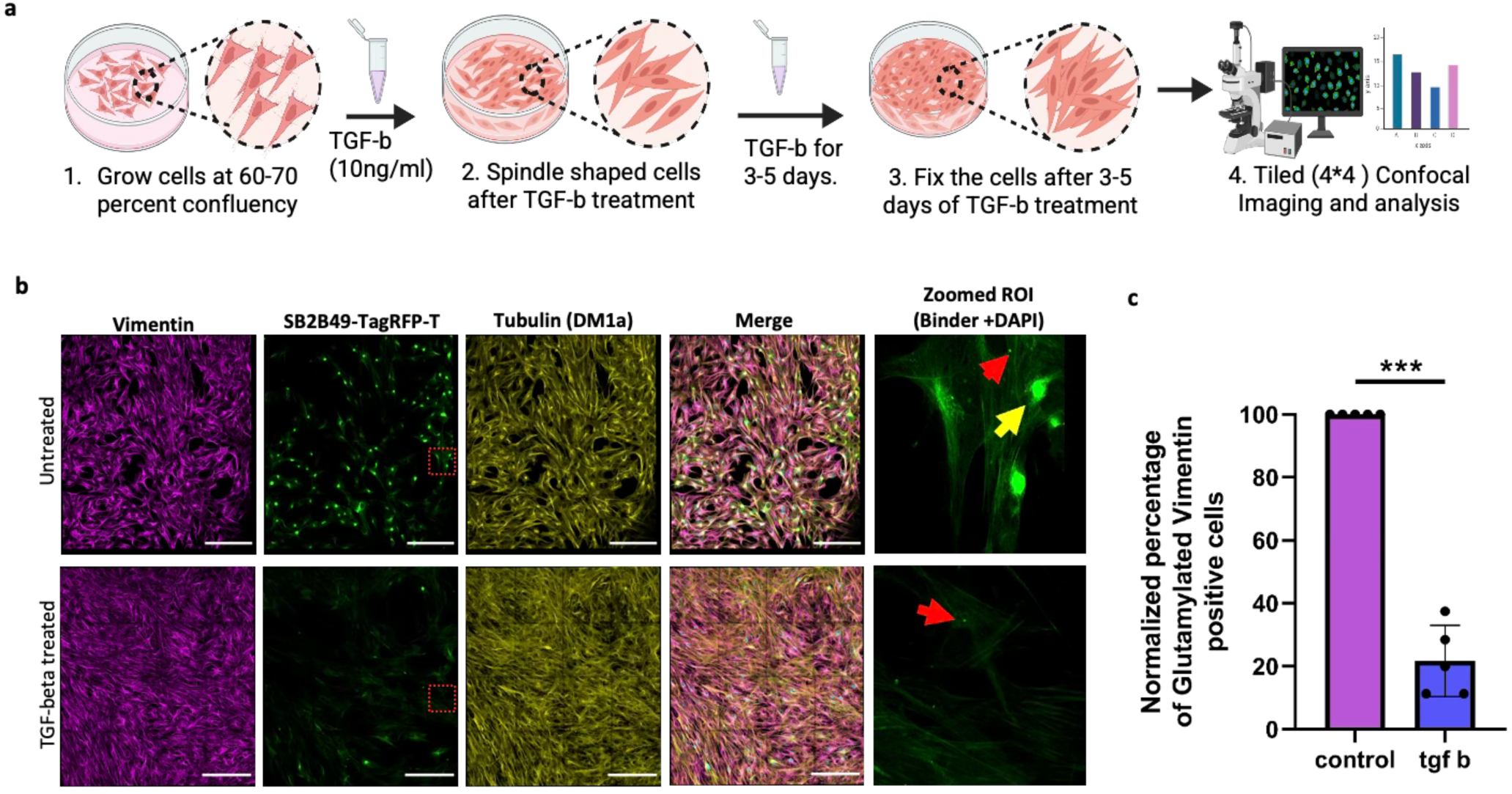
TGF-beta-induced EMT reduces glutamylated vimentin. **a.** Schematic representation of TGF-beta-induced epithelial to mesenchymal transition in RPE1 cells stably expressing the N-Tag RFPT fused SB2B49. **b.** Representative confocal tiled images of RPE1 cells stably expressing the N-Tag RFPT fused SB2B49 binder (green). The cells were continuously treated with TGF-beta for 5 days, fixed, and stained with B2-tubulin (yellow) and vimentin (magenta). Scale bar 200 μm. The dotted red box represents the zoomed-in region from control or TGF-beta-treated cells. The yellow arrow indicates cells with vimentin glutamylation, and the red arrows mark the centrosomes. **c.** Quantification from five different experiments representing the percentage of cells retaining the structure in TGF-beta-treated dishes, normalized to the control. Statistical significance was determined with Welch’s t-test. Each point represents an independent biological replicate. n=5. Error bars represent mean with SD, ***=P value <0.000.5.

During migration, cells transition from an epithelial to a mesenchymal state and upregulate vimentin, which plays a significant mechanical role and is often used as a marker for the amoeboid or mesenchymal state^42^. To determine whether glutamylated vimentin levels change during migration, we set up a cell migration assay (Fig. 6a). RPE1 cells stably expressing the glutamylation sensor were grown to confluence and scratched to create a wound (Methods). The cells were allowed to fill the gap between the two layers, and vimentin glutamylation levels were analyzed at different time points (Fig. 6a and Extended Data Fig. 9a-d). As cells fill the wound/scratch gap, the number of cells positive for glutamylated vimentin decreases relative to the distant field of view (Fig. 6b and Extended Data Fig. 9a-c). Post-scratch at the 18-hour time point, the wound/scratch gap is filled, and the cells regain the glutamylated vimentin modification (Fig. 6b, 6d, and Extended Data Fig. 9d). This suggests the glutamylated vimentin modification is dynamic and can be modulated during different cell states.

**Figure 6:**
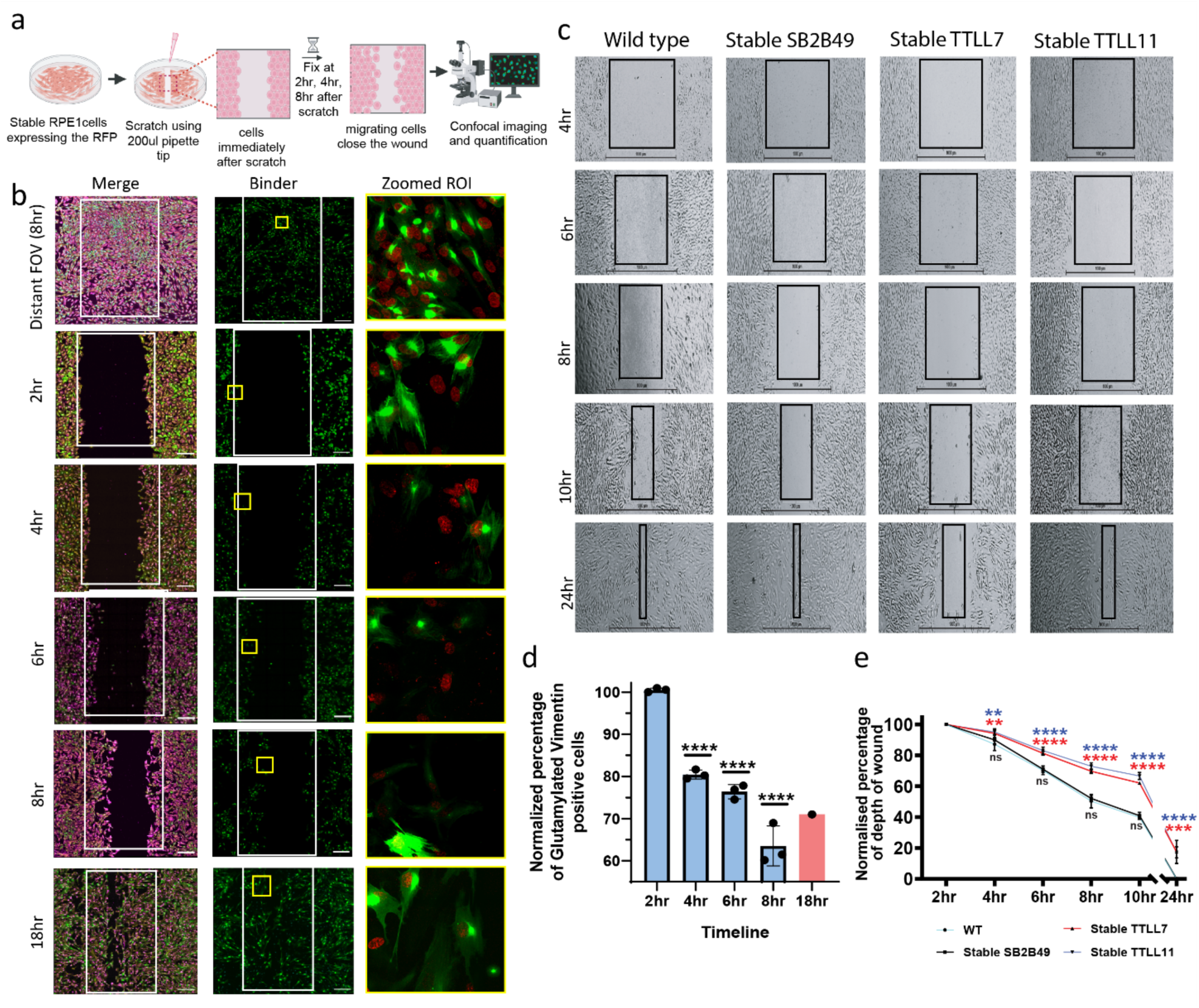
Wound-edge migration reduces glutamylated vimentin, and glutamylases delay wound closure. **a.** Schematic representation of scratch assay in RPE1 cells stably expressing the N-Tag RFPT fused SB2B49 (RPE1 high). **b.** Representative spinning disc microscopy tiled images of wounds in dishes with RPE1 cells stably expressing N-Tag RFPT fused SB2B49 binder (green). The Distant ROI represents the tiled image of a region far from the scratch. Scale bar 200 μm. The dotted yellow boxes represent the regions shown as zoomed in area from their respective tiled images. The cells were also stained with DAPI, Beta-2-tubulin, and vimentin, which are shown in red, yellow, and magenta, respectively. Only DAPI (red) and the N-Tag RFPT-fused SB2B49 binder (green) are shown in the zoomed-in ROI **c.** Representative brightfield images of RPE1 cells, wildtype (WT), stably expressing N-Tag RFPT fused SB2B49 binder (Stable SB2B49), TTLL7-eYFP (stable TTLL7), and TTLL11-eYFP (stable TTLL11), undergoing wound closure across different time points as indicated. Scale bar 1000μm. **d.** The graphical data representing the quantification of the cells retaining the structure (RPE1 high) over a period of 8hrs from the time of scratch. The data were normalized with control images of cells at a distant ROI (RPE1 high). Statistical significance was determined with Ordinary one-way ANOVA. Each point represents an independent biological replicate, n=3, ****= P value <0.0001 (see supplementary figure for all four channels) **e.** Percentage of wound closure area derived from experimental setup with WT, Stable SB2B49, TTLL7, and TTLL11 RPE1 cells across 3 different experiments. Error bars represent the mean with SD. Statistical significance was determined using a 2-way ANOVA. Each point represents an independent biological replicate, n=3, ****= P value <0.0001, ***= P value 0.0008, *= P value 0.0298. The color of the significance value corresponds to the color of the respective samples.

Next, we asked whether glutamylated vimentin modification influences cell migration. To address this, we stably expressed the glutamylases TTLL7 and TTLL11 in RPE1 cells and compared their migration rates with those of wild-type RPE1 cells and RPE1 cells stably expressing the glutamylation sensor (RPE1-Stable SB2B49) (Methods). While the wild-type and RPE1- Stable SB2B49 cells show comparable migration rates during wound closure, both TTLL7 and TTLL11 in RPE1 cells show decreased migration rates (Fig. 6c, 6e, and Extended Data Fig. 10a-d). These data collectively indicate an important role for glutamylated vimentin in cell migration and during the epithelial-mesenchymal transition.

## Discussion

Post-translational modifications can partition cytoskeletal polymers into functionally distinct pools, yet this principle is less well-defined for intermediate filaments than for actin filaments and microtubules^37^. Using multiple orthogonal approaches, we establish that a synthetic glutamylation binder (SB2B49) is a specific, glutamylation-dependent reporter of the glutamylated vimentin proteoform. The glutamylated vimentin proteoform offers a framework for reconciling cell-state transitions with vimentin biology. Vimentin is widely used as a mesenchymal marker at the protein level^16,19^, yet our findings emphasize the need to uncouple total vimentin from its proteoforms, especially during Epithelial-Mesenchymal Plasticity (EMP). Within epithelial-like contexts, our glutamylation sensor and binder delineate a subset of vimentin filaments rather than labeling the entire network.

Transcriptomic analyses of cells with glutamylated vimentin are enriched for epithelial programs. In addition to the epithelial-like state, our results show that the glutamylated vimentin proteoform is absent compared to total vimentin filaments in freshly thawed cells and emerges after a couple of passages (Extended Fig. 8b). While reviving can alter stress responses and enzyme expression, this passage-dependent appearance of glutamylated vimentin should be interpreted cautiously. However, these results further highlight the importance of distinguishing glutamylated vimentin levels from total vimentin filaments to more precisely categorize cellular states in both physiological and pathological conditions ^21–23,43^. During TGF-beta1-induced EMT, the glutamylation sensor positive pool is reduced, consistent with EMT-associated remodeling of the vimentin modification landscape. This suggests that EMP may involve not only changes in cytoskeletal expression and architecture, but also alterations in post-translational regulation and the reconfiguration of intermediate filaments. More broadly, these observations support the idea that intermediate filament networks, similar to microtubules, may be organized into functionally distinct substructures specified by post-translational modifications.

At the molecular level, regulation of glutamylated vimentin and its associated processes is mediated by the canonical glutamylation writer and eraser axes^3,44^. First, our overexpression experiments show that only TTLL6, TTLL7, and TTLL11 glutamylases increase the glutamylated vimentin pool, highlighting substrate selectivity within the TTLL family. Conversely, each of the tested cytosolic carboxypeptidases (CCPs) abolishes the glutamylated vimentin signal, suggesting their broad eraser capacity for this modification state. Second, increasing the glutamylation readout via overexpression of TTLL7 or TTLL11 is associated with decreased migration. Consistent with the possibility that glutamylation stabilizes a vimentin subnetwork that resists the extensive reorganization required during cell migration. While these experiments do not assign a unique writer–substrate pair, they establish that the glutamylation sensor and binder positive glutamylated vimentin pool is under active enzymatic control. This supports a model in which vimentin glutamylation is tuned by the balance of TTLL and CCP activities rather than emerging as a passive consequence of vimentin abundance. Phosphorylation of vimentin at Ser 56 and 71 has been previously reported and implicated in regulating vimentin filament dynamics and spatial organization^45,46^. Although we do not establish a direct causal link between vimentin glutamylation and filament dynamics, the dynamic behavior of glutamylated vimentin during cell migration and EMT suggests a central role. One parsimonious model is that glutamylation modulates the mechanical properties or interaction landscape of vimentin filaments, thereby shifting the balance between mechanical strength and malleability.

Distinguishing among these alternatives will require direct measurements of filament dynamics and mechanics coupled to defined perturbations of the modified sites.

A broader implication of our work is the development of a platform to generate synthetic binders that uncover enigmatic PTM-defined biology. These synthetic binders can directly enable visualization, enrichment, and functional dissection of modified proteoforms in their native cellular context. The systematic application of such binder-discovery platforms may accelerate the identification of new glutamylation substrates and reveal the context-specific roles of glutamylation across diverse cellular processes. SB2B49, the glutamylation sensor and binder described in this study, emerged from a screening platform originally developed against a glutamylated beta-tubulin carboxy-terminal tail epitope (B2-BiE). However, multiple orthogonal lines of evidence support SB2B49 as a reporter of glutamylation-dependent epitopes on both microtubules and vimentin filaments rather than a generic filament marker or charge sensor. An intriguing finding is the co-localization of the GT335 glutamylation-specific antibody and SB2B49 with vimentin filaments (Fig. 2b and 2e). So far, the GT335 antibody has been shown to recognize glutamylation branch points in both tubulin and non-tubulin glutamylation substrates^29,33^; however, there are no reports of vimentin glutamylation. Since SB2B49 was identified using a glutamylation branch point and shares similarities with GT335, including recognition of glutamylated vimentin, it is likely that SB2B49 recognizes the glutamylation branch point. Thus, the SB2B49 glutamylation binder described here can be applied to study context-dependent microtubule and vimentin glutamylation during various biological processes. In the case of the microtubule cytoskeleton, we have demonstrated its suitability for detecting *bona fide* microtubule glutamylation in cellular structures, including centrosomes and cilia basal bodies, mitotic spindle poles, the midbody during cytokinesis, axonemal and axonal microtubules (Figure 1 and Extended Fig. 3d-g). It is noteworthy that ectopic overexpression of glutamylase does not promote SB2B49 binding to cellular microtubules (Extended Data Fig. 4a and 4b), unlike conventional glutamylation-specific antibodies^32^.

Conversely, glutamylase overexpression promotes SB2B49 binding to vimentin (Extended Data Fig. 5a), suggesting that glutamylation sites are sparse and spread across the vimentin primary sequence, in contrast to the multiple-complex glutamylation molecular epitope landscape at the tubulin carboxy-terminal tail^37^. While mass spectrometry and mutational analyses support discrete glutamylation sites required for SB2B49 recognition, the stoichiometry and kinetics of site occupancy on vimentin remain unknown. Similarly, the SB2B49-positive pool robustly correlates with epithelial programs and is remodeled during EMT and migration; it is not clear whether vimentin glutamylation is necessary or sufficient to drive these transitions. Addressing these gaps will require targeted assays that directly connect defined modification states to filament dynamics, mechanics, and protein-protein interactions.

In summary, by coupling a generalizable synthetic binder discovery platform with biochemical validation and cell biology, our work provides a conceptual advance in understanding the partitioning of vimentin proteoforms during cell-state transitions. Our approach also provides a practical framework for discovering and interrogating glutamylation-dependent biology beyond tubulin. Given the tight coupling of vimentin glutamylation to epithelial programs and its dynamic loss during EMT and wound-edge migration, glutamylated vimentin may also serve as a state-sensitive biomarker. This can help stratify metastatic propensity or flag impaired wound-closure phenotypes in contexts where cytoskeletal remodeling is frequently dysregulated.

## Materials and Methods

### Antibodies and reagents

Alpha tubulin-DM1A Sigma-Aldrich Catalog no. T9026), , poly E (Adipogen, catalog no AG-25B-0030-C050), GT335 Sigma-Aldrich (Adipogen, catalog no AG-20B-0020-C100) , Vimentin (V9 (thermo fisher, catalog no MA5-11883) ,V1-10 (Thermo Fisher, catalog no MA1-10459), B2 abcam, catalog no ab15568), B3 (sigma, catalog no T2200), Tubulin tyrosiniated (sigma, catalog no T9028), Anti-Detyrosinated alpha Tubulin antibody (abcam, catalog no ab48389), Monoclonal Anti-tubulin acetylated (sigma, catalog no T6793), anti ARL13 b (Neuromab/UCDavis catalog no 75-287), anti-FLAG M2 (Sigma, catalog no F3165-0.2 mg). The anti-gamma-tubulin antibody was a kind gift from Dr. Sudarshan’s laboratory at the Institute for Stem Cell Science and Regenerative Medicine, India. DM1A, poly E, Anti-Detyrosinated alpha Tubulin antibody, and Monoclonal Anti-tubulin acetylated were all used at 1:3000-1:5000 dilution. Tubulin tyrosininated, anti-vimentin V1-10, V9, anti-FLAG M2, gamma tub, anti-ARL13 b, and anti-gamma tubulin were all used in 1:1000 dilutions unless stated otherwise. Secondary antibodies like Goat Anti Mouse Alexafluor-647 (catalog no A21235) Goat Anti-Rabbit alexafluor-647(catalog no A21245), Goat Anti Mouse Alexafluor-488 (catalog no A11001), Goat Anti-Rabbit Alexa-fluor 555 (catalog no A32732), Goat Anti Mouse alexa-568 (catalog no A11004), Goat Anti Mouse Alexafluor-647 (catalog no A21235), NucBlue live cell stain (catalog no r37605), DAPI (D1306) were all purchased from Invitrogen and used in 1:1000 dilution both in immunostaining and immunoblotting, Goat Anti-Mouse HRP (catalog no 626520) was purchased from Invitrogen and used in 1- 5000 to 1:7000 dilution. TGF b (R&D Systems, catalog no 240-B, retinoic acid, protein G agarose beads, PowerUP SYBR Green Master mix (catalog no A25742), Q5® Hot Start High-Fidelity 2X Master Mix (NEB, catalog no M0494L) and Quick-Load® Taq 2X Master Mix (NEB, catalog no M0271L)

### Peptide epitope designing and synthesis

The peptide of β2tubulin was designed featuring a bi-glutamylation modification on the amino acid residue (E435) that is found to be highly modified in the brain tissue and synthesized from LifeTein with a N terminus biotin tag. The peptide is named as B2-BiE, Biotin-DATADEQG-E-[(E(E)]-FEEEEGEDEA where -[(E(E)]- is the bi-glutamylation modification on E435 of GEF sequence in the peptide

### Screening of SSO7D yeast-display library

A combinatorial SSO7d yeast display library was obtained as a kind gift from Dr. Balaji M. Rao’s laboratory, and the detailed screening protocol for binders was adopted as described earlier^34,47^. The library, with a diversity of around 10^8^ unique clones, was propagated in SDCAA media (6.7 g/liter yeast nitrogen base without amino acids) [BD Difco; catalog no. 291940], 5 g/liter casamino acids [Gibco ; catalog no. 223050], 5.4 g/liter Na_2_HPO_4_ [Thermo-Fisher Scientific; catalog no. S374], 8.6 g/liter NaH_2_PO_4_.H_2_O [Merck; catalog no. 106346], and 1× penicillin-streptomycin [PenStrep; Gibco, catalog no. 15140122]). (20 g/liter D-(+)-glucose [Sigma-Aldrich; catalog no. G5767], was added prior to yeast inoculation, and the culture was grown at 30°C, 250 rpm for 18–24 h and passaged (3 to 4 times) until the expression exceeded 50 percent. The library was induced (10^9^ cells) in galactose-containing SGCAA media (20 g/liter D-(+)-galactose [Sigma-Aldrich; catalog no. G0750], 6.7 g/liter yeast nitrogen base, 5 g/liter casamino acids, 5.4 g/liter Na_2_HPO_4_, and 8.6 g/liter NaH_2_PO_4_.H_2_O) at 20°C for 20–24 h for the expression of binders on the surface of yeast cells.

#### Magnetic selection (MACS)

Around 10^9 nanobody-expressing yeast cells were pelleted and washed with PBS-BSA (8 g/liter NaCl, 0.2 g/liter KCl, 1.44 g/liter Na_2_HPO_4_, 0.24 g/liter KH_2_PO_4_, and 1 g/liter BSA). The cells were resuspended in 4.5 ml of PBS-BSA. Negative magnetic selection using streptavidin and anti-biotin microbeads was performed by incubating the yeast with 100 μl of anti-biotin microbeads (Miltenyi Biotec, catalog no. 130-090-485) and 100 μl of streptavidin microbeads (Miltenyi Biotec, catalog no. 130-048-102) at 4°C for 1 hour. The cells were pelleted, resuspended in 5ml PBS-BSA, and allowed to pass through (via gravity flow) the LD column (Miltenyi Biotec, catalog no. 130-042-901) placed on the Miltenyi MACS magnetic apparatus. The flow-through containing unbound cells was collected, and the column was washed with 2ml additional PBS-BSA to flush out any remaining cells. The cells were pelleted and resuspended in PBS-BSA for positive selection with the peptide epitope.

For positive selection, the negative-selected cells were incubated with 100 μM peptide at 4°C for 1 hour, followed by the addition of 100 μL anti-biotin microbeads and incubation for an additional 20 minutes at 4°C. The cells were pelleted, washed, and resuspended in PBS-BSA and passed through the pre-equilibrated LS column (Miltenyi Biotec, catalog no. 130-042-401) placed on the Miltenyi MACS magnetic apparatus. The unbound cells were washed off with PBS-BSA, and the bound cells were isolated from the column using a 50ml Falcon. The cells were pelleted, resuspended in glucose-containing SDCAA media, incubated at 30°C and 250 rpm for 48 hours, and then expanded further.

#### Fluorescent Activated Cell Sorting (FACS)

The MACS-sorted culture was induced (around 10^9 cells) in galactose media at 20°C for 72 hours. Around 10^7 cells were taken in a 1.5ml microcentrifuge tube, washed with 1ml PBS-BSA, and resuspended in 100 μl PBS-BSA. The cells were incubated with 100 µM of Biotinylated B2-BiE peptide. and 1:200 dilution of rabbit anti-HA tag antibody (Sigma; catalog no. Cat # H6908) at 4°C for 1hr. 30minCells were pelleted, washed three times with PBS-BSA, and incubated with a 1:200 dilution of goat anti-rabbit Alexa fluor-647 antibody (Invitrogen) and a 1:100 dilution of neutravidin fluorescein conjugate (Invitrogen; FITC, catalog no. A2662). Cells were resuspended in 100 μL PBS-BSA and sorted on BD FACS Aria Fusion for double-positive cells, with proper controls including unstained and single-stained ones. Around 3000-5000 double-positive cells were collected, grown in 5 ml fresh SDCAA media, and propagated in larger volumes of SDCAA media (250–500 ml). The FACS was repeated 5 times with the decreasing peptide concentration of 100uM, 75uM, 50uM, 50uM, and 25uM in each subsequent FACS round. Stocks were made after each FACS round. After the fifth round of sorting, the cells were grown, expanded, and plated on SDCAA agar (20 g/liter dextrose, 6.7 g/liter yeast nitrogen base, 5 g/liter casamino acids, 5.4 g/liter Na_2_HPO_4_, 8.6 g/liter NaH_2_PO_4_ · H_2_O, 182 g/liter sorbitol, and 15 g/liter agar).

### Sanger sequencing

The 44 SS07D sequences were amplified directly from the yeast colonies using the following primers:

pctcon_ss07d FP:

5’-gtcaacgactactattttggccaacgggaaggc-3’

pctcon_ss07d RP:

5’-gctatataaagtatgtgtaaagttggtaacggaacg-3’

The sequences were subsequently subjected to Sanger sequencing using the following primers.

pctcon_ss07d seq FP:

5’-tacgacgttccagactacgctct-3’

pctcon_ss07d seq RP

5’-gggaaaacatgttgtttacggag-3’

The sequences were analysed using Snapgene^TM^ and the multiple sequence alignment was done using Clustal Omega.

### Cloning for bacterial expression

The 32 unique sequences were cloned in pet28a with a mCherry Tag at C-terminus using the following primers

Forward primer:

5’-cacagcagcggcctggtgccgcgcggcagccatatggctagcatggcgaccgtgaaatttaaatataaaggcg-3’

Reverse primer:

5’-ggccatgttatcctcctcgcccttgctcaccatagcgctgccaagcttttttttctgtttttccagcatctgcagc-3’

SB2B49 was cloned in pet28a vector with a KCK motif at its C terminus for maleimide labelling with Alexa fluor dyes using the following primers:

Forward primer:

5’-gtttaactttaagaaggagatataccatggatgcatcatcatcatcatcacagcagcggcgaaaacctgtattttcagggcggatccatggcgaccgtgaaatttaaatataaaggc-3’

Reverse primer:

5’-ctcgagtgcggccgcaagcttgtcgactcatttgcattttccggattttttctgtttttccagcatctgcag-3’

### Cloning for mammalian expression

All cloning was done in the lentiviral pTRIP vector under the CMV enhancer and the chicken β-actin promoter (CAG promoter), flanked by 5 ′ and 3′ long terminal repeat sequences. SB2B49 was cloned with N-terminus Tag RFPT employing three fragment Gibson assembly cloning using the following set of two primer pairs to amplify SB2B49 and Tag RFPT, respectively. This vector was named as pTRIP-N Tag RFPT-SB2B49.

Forward primer 1:

5’-catcattttggcaaagaattattccgctagccaccatggtgtctaagggcgaagagctgattaaggagaacatgc-3’

Reverse primer 1:

5’-cctttatatttaaatttcacggtcgccactccggaggatcccttgtacagctcgtccatgcc-3’

Forward primer 2:

5’-ggcatggacgagctgtacaagggatcctccggagtggcgaccgtgaaatttaaatataaagg-3’

Reverse primer 2:

5’-gctccatgtttttctaggtctcgaggtcgactcattttttctgtttttccagcatctgcagc-3’

SB2B49 was cloned along with a FLAG tag at the C-terminus by digesting the above vector using BamHI and SalI restriction sites and amplifying the SB2B49 using the following primers

Forward primer:

5’-ggcatggacgagctgtacaagggatcctccggagtggcgaccgtgaaatttaaatataaagg-3’

Reverse primer:

5’-catgtttttctaggtctcgaggtcgacttacttgtcatcgtcgtccttgtagtcgctaccttttttctgtttttccagcatctgcagc-3’

Since the SS07D is a bacterial protein, to increase the efficiency of expression of SB2B49 in mammalian cells, it was codon-optimized from Twist Biosciences and cloned into the pTRIP vector by digesting the pTRIP-N Tag RFPT-SB2B49 with BamHI and SalI restriction sites and amplifying the Opti SB2B49 using the following primers

Forward primer:

5’-ggcatggacgagctgtacaagggatcctccggagtcgccacagttaagttcaagtacaag-3’

Reverse primer:

5’-gctccatgtttttctaggtctcgaggtcgactcacttcttttgcttctcaagcatttgtaacaattc-3’

Vimentin was cloned into the pTRIP vector along with a C-terminal eGFP tag using three fragment Gibson assembly cloning using the following primer pairs to amplify vimentin and eGFP, respectively.

Forward primer 1:

5’-CTcatcattttggcaaagaattattccgctagcatgtccaccaggtccgtgtcctcgtcctccTACCGC-3’

Reverse primer 1:

5’-Cagctcctcgcccttgctcaccatgtcgactgaaccggatccttcaaggtcatcgtgatgctgagaag-3’

Forward primer 2:

5’-Cttctcagcatcacgatgaccttgaaggatccggttcagtcgacatggtgagcaagggcgaggagctg-3’

Reverse primer 2:

5’-gattgctccatgtttttctaggtctcgagcggccgctttacttgtacagctcgtccatgccgagagtg-3’

Vimentin was gene-synthesized by Twist Biosciences with nine stretches of glutamate-to-aspartate mutations. The synthesized vimentin gene with mutations was then cloned in pTRIP vector by replacing WT vimentin with a C-terminal eGFP tag. The following primers were used to amplify the Mutant vimentin sequence, which was then cloned using two-fragment Gibson cloning.

Forward primer:

5’-Cttctcagcatcacgatgaccttgaaggatccggttcagtcgacatggtgagcaagggcgaggagctg-3’

Reverse primer:

5’-gattgctccatgtttttctaggtctcgagcggccgctttacttgtacagctcgtccatgccgagagtg-3’

In addition to the above-mentioned mutations, the following site-directed mutagenesis (SDM) primers were designed to mutate a single site in addition to the above-mentioned primers at either E95 or E103.

Forward primer E95D :

5’-gacgccatcaacaccgacttcaagaacacccgcaccaa-3

Reverse primer E95D :

5’-ttggtgcgggtgttcttgaagtcggtgttgatggcgtc-3 Forward primer E103D :

5’- ttcaagaacacccgcaccaacgacaaggtggagctgcaggag -3

Reverse primer E103D :

5’- ctcctgcagctccaccttgtcgttggtgcgggtgttcttgaa -3

### Protein purification

All the SB2B49 or SB2B49 with mCherry or KCK motif were purified from Rosetta (DE3)–competent cells. The pet28a plasmids carrying the gene of interest were transformed into Rosetta (DE3)–competent cells, and the resulting bacterial colonies were inoculated into 10ml LB medium and incubated overnight at 37°C and 180 rpm. The next day, the cultures were inoculated in larger volumes of LB medium and induced at an OD600 of 0.5 with 0.5 mM IPTG (Sigma-Aldrich; catalog no. I6758) at 20°C, overnight. The cells were harvested in 50 mM HEPES, pH 7.5, 100 mM KCl, and 1 mM PMSF, with 1 tablet of EDTA-free protease inhibitor cocktail (Roche; catalog no. 11836170001) for 1 liter of culture, and additionally with 1 mM TCEP for KCK containing SB2B49. The cells were lysed with the Ultrasonic homogenizer (SJIA LAB, SKL-500D) or Avestin Emulsiflex C3 homogenizer (ATA Scientific Instruments), The lysate was centrifuged and the supernatant loaded onto a 5 ml Ni-NTA affinity column, washed with a 10–column volume wash with 50 mM HEPES, pH 7.5, 500 mM KCl, containing either 25 mM imidazole and 50mM imidazole, pH 7.5, and eluted in 5 column volumes of 50 mM HEPES, pH 7.5, 100 mM KCL, and 500 mM imidazole, pH 7.5). The eluted protein was concentrated up to 5 ml using a 3-kD Millipore Amicon filter (Merck; UFC900324). Further, size exclusion chromatography was performed in 50 mM HEPES, pH 7.5, and 100 mM KCL buffer with an additional 1mM TCEP for the KCK motif containing SB2B49 in a Superdex-75, 16/600 column (Cytiva; catalog no. 28989333).

### Maleimide labelling of the SB2B49 with CF640R

Purified KCK motif containing SB2B49 was mixed with the CF^®^640R (biotium, catalog no. 92034) dye in the molar ratio of 1:10, with a final concentration of 100uM and 1mM of the protein and dye, respectively, with the volume not exceeding more than 180ul. The mixture was then incubated at 4 °C, with the rotor running overnight. The next day, the mixture was added to the PD SpinTrap G-25 column (Cytiva, catalog no. 28918004 ) as per the manufacturer’s instructions. Briefly, after removing the bottom closure, the column was equilibrated with buffer containing 50 mM HEPES (pH 7.5), 100 mM KCl, and 1 mM TCEP. Around 180 μL of the protein-dye mixture was applied to the column in the middle of the packed bed. The protein was then eluted by centrifuging the column at 800g for 2min. The labelling efficiency was measured by diluting the protein in 8M guanidine hydrochloride, and the absorbance at 280nm and 647nm was measured. The labelling efficiency was calculated using the following formula:

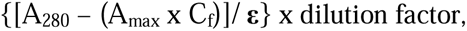

where C_f_ is the dye correction factor, ε is the coefficient of extinction, A_280_ and A_max_ are the absorbance of the protein dye conjugate at 280nm and the absorption maximum of the dye, respectively.

### In-vitro lysate-based microtubule binding assay for binders

Following sequence verification, the clones were transformed into Rosetta (DE3)–competent cells, and the resulting bacterial colonies were inoculated into a deep 96-well plate containing LB medium with 50 μg/ml kanamycin and chloramphenicol, with 1 ml of medium per well. The cells were grown overnight in LB medium at 37 °C, 180rpm. The next morning, 10 μL from each well was inoculated in a fresh deep 96-well plate containing LB medium and grown at 37 °C, 180rpm until the OD_600_ of the cultures reached around 0.5 -1. The OD_600_ of random wells was measured and averaged at 0.5-1 (2-3hrs). The cultures in the plate were at OD_600_ of 0.5 with 0.5 mM IPTG (Sigma-Aldrich; catalog no. I6758) at 25°C overnight. The plate was centrifuged at 1000 rpm for 10 min at 4°C, and the pellets in each well were washed with 50 mM HEPES, pH 7.5, 100 mM NaCl, then resuspended in 200 μL of the same buffer and sonicated for 1 min per well. The plate was centrifuged and the supernatant transferred to a fresh 96-well plate, which was either stored in -80°C or used for microtubule binding TIRFM assay. To crudely determine the affinity of binders towards modified microtubules, microtubules were polymerised using tubulin isolated from goat brain in the presence of 2 mM GTP (Sigma-Aldrich). and 20 µM Taxol (Sigma-Aldrich) at 37°C in 1× BRB80 buffer (80 mM PIPES, 1 mM MgCl_2_, and 1 mM EGTA, pH 6.8, with KOH). 10-20 µM unlabelled tubulin was set for polymerization along with Alexa Fluor 647–labeled tubulin (1/10 molar ratio), A glass chamber made up with coverslip glass (0.17 mm thick) was coated with rigor kinesin, followed by a 1× BRB80 wash and the surface was blocked 1.25 mg/ml β-casein (Sigma-Aldrich; catalog no. C6905) in 1× BRB80 buffer. 10–20-fold diluted polymerized microtubules were flowed into the flow chamber for 3min and subsequently washed with 1× BRB80 containing 1.25 mg/ml β-casein and 20 µM Taxol. Before the addition of the binder lysates, the chambers were washed with 50 mM HEPES pH 7.5, 100 mM NaCl. The binder lysates from each well of the 96-well plate were added to the flow chambers in such a way that a new binder lysate was added after the previous binder had been imaged, and no two binders were added to the same chamber. Microtubules were first imaged using a Nikon Eclipse Ti-2 H-TIRF microscope with the red laser (640 nm) for Cy5-labelled microtubules and the green laser (561 nm) to visualize the microtubule-bound binders.

Similarly, to determine the specificity of the binders towards modified (brain microtubules) vs unmodified (HeLa microtubules), HeLa microtubules were prepared and polymerized as described above without any labeling. Both types of microtubules were added to the flow chamber, and the HeLa microtubules were visualised using GFP-tagged A1AY1 (tyrosination sensor, Kesarwani et al 2020. The binder lysates were added to the flow chambers and imaged using a Nikon H-TIRF microscope.

### Cell culture

The RPE1 cell line was a kind gift from Dr. Sudarshan’s laboratory at the Institute for Stem Cell Science and Regenerative Medicine, India. MDA MB231 and MCF 7 were a kind gift from Prof Apurva Sarin’s laboratory, Institute for Stem Cell Science and Regenerative Medicine, India. The SH-SY5Y cell line was a kind gift from Prof. Gaiti’s laboratory at NCBS, India. RPE1 and SH-SY5Y cells were cultured in DMEM/F12 medium (Gibco, Catalog no 10565018) supplemented with 10% FBS and 1x Pen-Strep. HEK 293[T, NIH 3T3, MDA MB231, MCF 7 cells were cultured in DMEM containing 10% FBS, supplemented with 1× sodium pyruvate (1 mM; Gibco), 1× GlutaMAX (Gibco). U2OS cells were grown in McCoy’s 5A (Sigma-Aldrich; M4892) medium supplemented with 10% FBS and 1x Pen-Strep. RPE 1 cells in either 35 mm Ibidi dishes or 8-well slides were treated with 10ng/ml Recombinant Human TGF-beta 1 Protein (R&D Systems, catalog no 240-B) for 3-5 days as specified in the legends. Stable SH-SY5Y expressing SB2B49 with an N Tag RFPT was grown to 50 % confluency, after which it was incubated in medium without FBS, supplemented with 10 μM retinoic acid (Merck, Catalog no. R2625-1G), for 3-5 days, with fresh retinoic acid added each day. The cells were fixed and imaged as mentioned in the methods.

### Transient transfection protocol for imaging and western blot experiments

All transient transfections in mammalian cells, including HEK, RPE1, and U2OS, were performed at ∼60–70% cell confluency in 35-mm Ibidi dishes (glass-bottom coverslip surface Ibidi; catalog no. 81218) or 8 Well slides (µ-Slide 8 Well high Glass Bottom, catalog no 80807) using Lipofectamine 3000 as per the manufacturer’s protocol. Briefly, 200 ng or 2 µg of plasmid was used for each well in an 8-well slide and a 35mm dish, respectively. The DNA was added to 5 μL or 125 μL Opti-MEM, and 0.5 or 2 μL of P3000™ Reagent for 8-well and 35 mm dishes, respectively. In another tube, Lipofectamine™ 3000 Reagent was added in a 1:1 (V/V) ratio to an equal amount of Opti-MEM, following which both tubes were combined, and the mixture was incubated for 10 - 20 min at room temperature (RT). The mixture was added to dishes containing freshly prepared 10% serum–containing media and incubated for either 6 hours or overnight. The next day, the cells were stained with 0.5–1 µM SiR-tubulin (Cytoskeleton; catalog no. CY-SC002 Spirochrome kit) for 1 h before imaging. Cells were imaged on an FV3000 Olympus confocal microscope after 24 h of transfection. Similarly, for co-transfection experiments, the DNA concentration per dish was not exceeded compared to that used for single transfections. All the vimentin constructs were transfected into 8-well slides For Western blots or pull-down experiments, either 100 mm or T75 dishes were used. The concentration of DNA transfected did not exceed 10 μg, and lipofectamine reagents were used at a 1:1 V/V concentration.

### Generation of stable cell lines

HEK293-T cells were cultured until about 70-80% confluency in complete DMEM media containing 10% FBS, 1× GlutaMAX (Gibco), 1× sodium pyruvate, and 1× PenStrep in a humidified 37°C incubator with 5% CO_2_. Lentiviral production in HEK293-T cells was performed using pTRIP vector constructs for all genes. For a 100-mm or T75 dish transfection, 5 µg of lentiviral plasmid with the gene of interest, 3.75 µg of psPAX2 (Addgene; #12260), and 1.25 µg of pmDG2 (Addgene; #12259) plasmid were mixed in 500 µl of OptiMEM media with 20 µl of P3000 (Invitrogen; Lipofectamine-3000, catalog no. L300015) or 10 µl of PLUS reagent (Invitrogen; LTX, catalog no. L15338100). Separately, 500 µl of OptiMEM was combined with 30 µl of Lipofectamine-3000 or Lipofectamine-LTX reagent. The tubes were incubated separately for 5–10 minutes at room temperature, then combined and incubated for another 15 minutes before being added dropwise to the dishes. The cells were maintained in a 37°C incubator with 5% CO_2_ for approximately 13–15 hours, after which the transfection media was replaced with DMEM containing 10% FBS. Media was collected at 24 and 48-hour intervals and concentrated using a 50-kD Millipore Amicon filter (Merck; UFC905024) at 1,000g to approximately 1–3 ml. The concentrate was then supplemented with Lenti-X concentrator (Takara; catalog no. 631231) at a 1:3 volume ratio and incubated overnight at 4°C. The next day, the mixture was pelleted at 1,500g for 45 minutes at 4°C, resuspended in 1 ml of DMEM with 10% FBS, and stored at −80°C for long-term use. Lentiviral transduction was performed on 60% confluent cells in a 24-well plate with varying viral titers, in the presence of 1 µg/ml of polybrene (Merck; catalog no. TR-1003-G). After 24 hours, the cells were passaged and expanded as usual.

### Immunostaining of mammalian cells and mouse sperm with Alexa Fluor 647 labelled or mCherry tagged SB2B49

The RPE1 cells were grown in either 8-well or 35mm Ibidi glass-bottom dishes until 60-70% confluency and either fixed with 4% paraformaldehyde or first transfected with eYFP-tagged TTLL enzyme plasmids and then fixed. Cells were fixed after at least 24 h from seeding. Cells were rinsed multiple times with 1× BRB80 and fixed with 4 percent paraformaldehyde in BRB80 for 15min at RT. Cells were washed three times for 5 min each, then permeabilized with 0.1% Triton-X-100 in 1× BRB80 for 10 min at room temperature. Cells were blocked for 1 h at room temperature with 5% BSA made in 1× BRB80. Cells were incubated overnight at 4°C or for 2 hours at RT with the primary antibody. Cells were washed three times for 10 min each at room temperature and incubated with the fluorescently tagged secondary antibody for 2 h at room temperature or overnight at 4 °C. The cells were washed with buffer containing 50 mM HEPES (pH 7.5), 100 mM KCl, 1 mM TCEP, and 5% BSA. The binder was added to the cells at concentrations of 1 μM to 5 μM, and the cells were incubated overnight at 4°C in the same buffer. The next day, the cells were washed three times before imaging on the Olympus FV3000 confocal microscope. Sperm was isolated from mice kindly provided by Dr. Sudarshan, permeabilized with 1X BRB80 containing 0.01% Triton-X 100 for 5 min, and then fixed with 4% paraformaldehyde. The sperm were fixed in 5% BSA in 1x BRB80 for 1 hour at RT, after which the protocol described above was followed.

### Immunoprecipitation assay

RPE1 cells expressing the binder (RPE1 high) were pelleted from a T75 flask (about 3 x 10^6 cells). The cells were lysed with ice-cold 50 mM HEPES, 100 mM KCl, pH 7.5, 0.2% Triton X-100 buffer for 1 h. Next, the cells were sonicated for 1 minute, and the lysates were centrifuged at 13000 rpm for 30 minutes. 500-700 μg of supernatant was incubated with Protein A Dynabeads (Invitrogen, catalog no. 10002D) overnight at 4°C. The beads were pre-incubated with 1:500 Anti-FLAG M2 antibody (Sigma, F3165-0.2 mg) for 6 hours at 4°C. The bead- antibody-binder complex was washed 5- 6 times with 50[mM HEPES, 100mM KCl, pH 7.5, 0.5% Triton X-100, followed by elution in 75 μL of TGS containing 1× Laemmli buffer (180 mM DTT [Sigma-Aldrich; #D9779], 4% SDS [VWR; #442444H], 160 mM Tris-HCl, pH 6.8, 20% glycerol [VWR; #24388.295], and bromophenol blue). The mixture was boiled at 95°C for 10 minutes. The supernatant was collected into a fresh tube and loaded onto an SDS-PAGE gel for western blotting or mass spectrometry. 10-20 µL and 5-10 µL of the eluate were used for SDS-PAGE in western blot and mass spectrometry, respectively.

### Migration assay

For the wound-healing assay, RPE1 cells were seeded into multiple 35mm Ibidi dishes and grown to 100% confluency. Cells were washed with 1x DPBS. A scratch was given using a 200 ul pipette tip, and the dishes were washed with 1x DPBS. Fresh medium was added, and the dishes were fixed at 2, 4, 6, and 8 hrs post-scratch, as described above, and imaged. For the TTLL-related migration assay, two dishes were seeded per condition, and a scratch was created using a 200ul tip. Brightfield images were taken at 2, 4, 6, 8, 10, and 24 hrs post-scratch using a Nikon Eclipse TS-2 microscope with a 4x objective.

### Image acquisition and analysis, graphs and statistical analysis

All the images were acquired using an inverted confocal microscope (Olympus FV3000) equipped with solid-state laser lines (405, 488, 561, and 640 nm) and a 60× oil objective unless stated otherwise. All images were taken with ibidi glass-bottom dishes. To image a larger area of interest in the TGF-beta1–induced EMT experiment, RPE1 cells were first imaged with a 10× objective to capture a broad view. This was then used to map the fields of view in the Olympus FluoView™ software as a 4×4 stitched image, with each field covering 1024[×[1024 pixels. Imaging of cells in the wound-healing assay was performed using a Nikon spinning-disk confocal microscope. RPE1 cells were imaged with a 10× objective to encompass the entire scratch area, and this image was used as a preview in the xyz overview plugin in the NIS-Elements AR software. The final image was generated by stitching 8×8 fields of view, each with a 2048×2048-pixel resolution. An equivalent area far from the scratch was also imaged for analysis.

All images were analyzed using Fiji (ImageJ) software, and the images in the manuscript are maximum-intensity projections (MIPs) of Z-stacks. Cell counting for the TGF β experiment and wound healing assay was performed using the Cell Counter plugin in Fiji, with nuclei used as a reference for the total number of cells counted. The Cell Counter plugin was used to count the number of glutamylated vimentin-positive cells relative to the total number of enzyme-positive cells (Fig. 2) and vimentin eGFP-positive cells (Fig. 3). All cells were manually counted, and the binder channel type was set to 2. For the colormap shown in Fig. 2, the Fiji plugin “colocalization colormap” was used to obtain the colocalization value (index of correlation, icorr) and the colormap distribution. A 50–100-pixel background was subtracted from the antibody and Binder channels.

All graphs and statistical analyses were generated using the commercially available GraphPad Prism software. Statistical analysis performed for mean ratio, SD, and n values is mentioned in the respective figure legends.

### Transcriptomic analysis

The preliminary quality control (QC) was checked using FASTQC (Version 0.12.1) before and after trimming the adapters. The adapters were trimmed using Trimmomatic (Version 0.39). Alignment QC was performed using the alignment statistics obtained from STAR (Version 2.7.3a) alignment with the Hg38 reference sequence. The raw read counts were estimated using FeatureCount (Version 2.24.0). Read count data was normalized using DESeq2 (Version 1.50.2). The ratio of normalized read count for test over control was taken as the fold change. Genes were first filtered based on the p-adjusted value (<0.05). Those genes that were found to have ±1 log2 (fold change) were considered as statistically significant. For samples without replicates, fold change was calculated using NOISeq (Version 2.54.0). Genes with a q-value equal to 0.8 were considered statistically significant. GO annotation and Reactome pathway information for differentially expressed genes were done using clusterProfiler (Version 4.18.2) mentioned in the respective figure legends.

### Mass spectrometry and data analysis Protein digestion and Peptide extraction

Protein Immunoprecipitation samples were subjected to run on 10% SDS polyacrylamide gel up to 1cm in length (into the resolving gel). The SDS PAGE was stained using Coomassie brilliant blue R-250 and destained with SDS PAGE destain solution with 50% Water, 10% Acetic acid, 40% Methanol. The protein band was cropped out of the gel with a surgical blade and further chopped into small pieces of approximately 2 to 4 mm. The gel pieces were washed further with Mass Spectrometry-grade water three times and then destained with 50% Acetonitrile in 100mM Ammonium Bicarbonate until the gel pieces become transparent. Protein digestion was performed on in-gel samples using mass-grade Trypsin as per the standard protocol. In brief, gel pieces were dehydrated with Acetonitrile and destained, and then dried in a speed vac. Samples were then subjected to digestion with 200ng of Trypsin in ammonium bicarbonate buffer for overnight at 37 0 C. After the digestion, the peptides were extracted from the gel pieces by adding 50%

Acetonitrile and 0.1% Formic acid in triplicate. Extracted peptides were pooled together and dried in a SpeedVac. Peptides were reconstituted in 2% Acetonitrile and 0.1% Formic acid (20µL) before the injection of the peptide into the mass spec for analysis.

### LC-MS analysis of digested peptides Separation of peptides on Micro LC

Peptides were reconstituted, and 2 µL of the peptides were injected into the Sciex M5 Micro LC in trap elute mode. The peptide was desalted online for 5 min at a 10 µL flow rate on nanoEase TM M/Z HSS T3, 100 A 0, 5 µm, 300 µm x50mm Trap column (Waters) and then the desalted peptides were subjected to separation on nanoEaseTM M/Z HSS T3, 100 A 0, 1.8 µm, 300 µm x100mm analytical column (Waters) with flow rate of 5 µL/min. Peptides were resolved with the gradient concentration of two mobile phases, Mobile phase B ( 95% Acetonitrile, 0.1% Formic acid) and mobile phase A (5% Acetonitrile and 0.1% Formic Acid) for 17 min on an analytical column with flowrate of 5 µL/min. In brief, the mobile phase gradient concentration was changed from A to B as per the following table.

| Time | % Mobile Phase-B | Mobile phase-A | Flow rate (µL/min) |
| --- | --- | --- | --- |
| 0 | 5 | 5 | 5 |
| 2 | 10 | 90 | 5 |
| 10 | 30 | 70 | 5 |
| 11 | 95 | 5 | 5 |
| 13 | 95 | 5 | 5 |
| 14 | 5 | 95 | 5 |
| 17 | 5 | 95 | 5 |

### Mass spec data acquisition of Peptides extract

Sciex M5 Micro LC was connected to the Sciex High Resolution 7600 Zeno TOF mass spec for data acquisition. Sciex Zeno TOF 7600 is configured with an Opti Micro flow ESI source for better sensitivity. The mass spectrometry data were acquired using Sciex OS 3.2. The TOF MS data were acquired in the mass range of 350 Da to 1250 Da, whereas the MS/MS (fragment ion) data were acquired in the mass range of 150 to 1500 Daltons. During the data acquisition, the dynamic collision energy was used for the fragmentation of the peptide. Zeno trapping of 0.02msec was used to increase the sensitivity of the peptide fragment, whereas the 20 most abundant peptide was fragmented in each cycle. Only TOF MS peaks having 2 or more than 2 to 5 charge states were fragmented for peptides.

### Database search

Mass spec data for the Immuno pull down samples were searched using Proteome discoverer 2.0 and Uniprot Human Proteome Database using semi-specific trypsin cleavage and Cys=57.02146 Da as static modification whereas dynamic modification was set for Met=15.99492 Da, Mass spec data for the Immuno pull down samples were searched using Proteome discoverer 2.0 and Uniprot Human Proteome Database using semi-specific trypsin cleavage and Cys=57.02146 Da as static modification whereas dynamic modification was set for Met=15.99492 Da, Unimod accession no. “451,452 and 453” Delta mass 258.08519, 387.12778, and 516.17037 for GluGlu, GluGluGlu and GluGluGluGlu. Results were identified with Percolator q-value < 0.001.

## Acknowledgements

We are grateful to the Central Imaging and Flow Facility (CIFF) and the Mass Spectrometry Facility at the iBRIC-inStem and Bangalore Life Science Cluster, India. M.S acknowledges funding support from iBRIC-inStem core grants from the Department of Biotechnology, India, DBT/Wellcome Trust India Alliance Senior Fellowship (IA/S/22/2/506502), EMBO Young Investigator Programme award, and ANRF-CRG grant (CRG/2023/005854) from the Department of Science and Technology (DST), India. S.G is supported by the iBRIC-inStem core grants from the Department of Biotechnology, India, DBT/Wellcome Trust India Alliance Intermediate Fellowship (IA/I/22/1/506238) and ANRF start-up research grant (SRG/2023/000847) from the DST, India. R.G is supported by a CSIR-SRF. I.K, A.B acknowledge funding support from iBRIC-inStem core grants from the Department of Biotechnology, India. L.P.D and S.M are supported by the DBT-JRF.

## Author Contributions

R.G and M.S conceptualized the project. R.G screened, validated, and performed the biochemical and microscopy experiments with the glutamylation sensor and binder. A.B, S.S, I.K, and L.P.D cloned the binder and vimentin clones and purified the recombinant glutamylation binder labelled with CF640R. S.G and S.M provided the HEK tubulin, mouse sperm, and TTLL11 constructs. G.S analyzed the RNA transcriptomic data. N.S. performed the mass spectrometry experiments and analyses and provided insights into the data. M.S. supervised and secured funding for the project. R.G and M.S interpreted the results and wrote the manuscript.

### Disclosure and competing interests statement

M.S. and R.G are inventors on a provisional patent application describing the invention and application of the Glutamylation sensor, binder sequences, and glutamylated vimentin epitopes described in this manuscript.

---

**Extended Data Fig 1.**
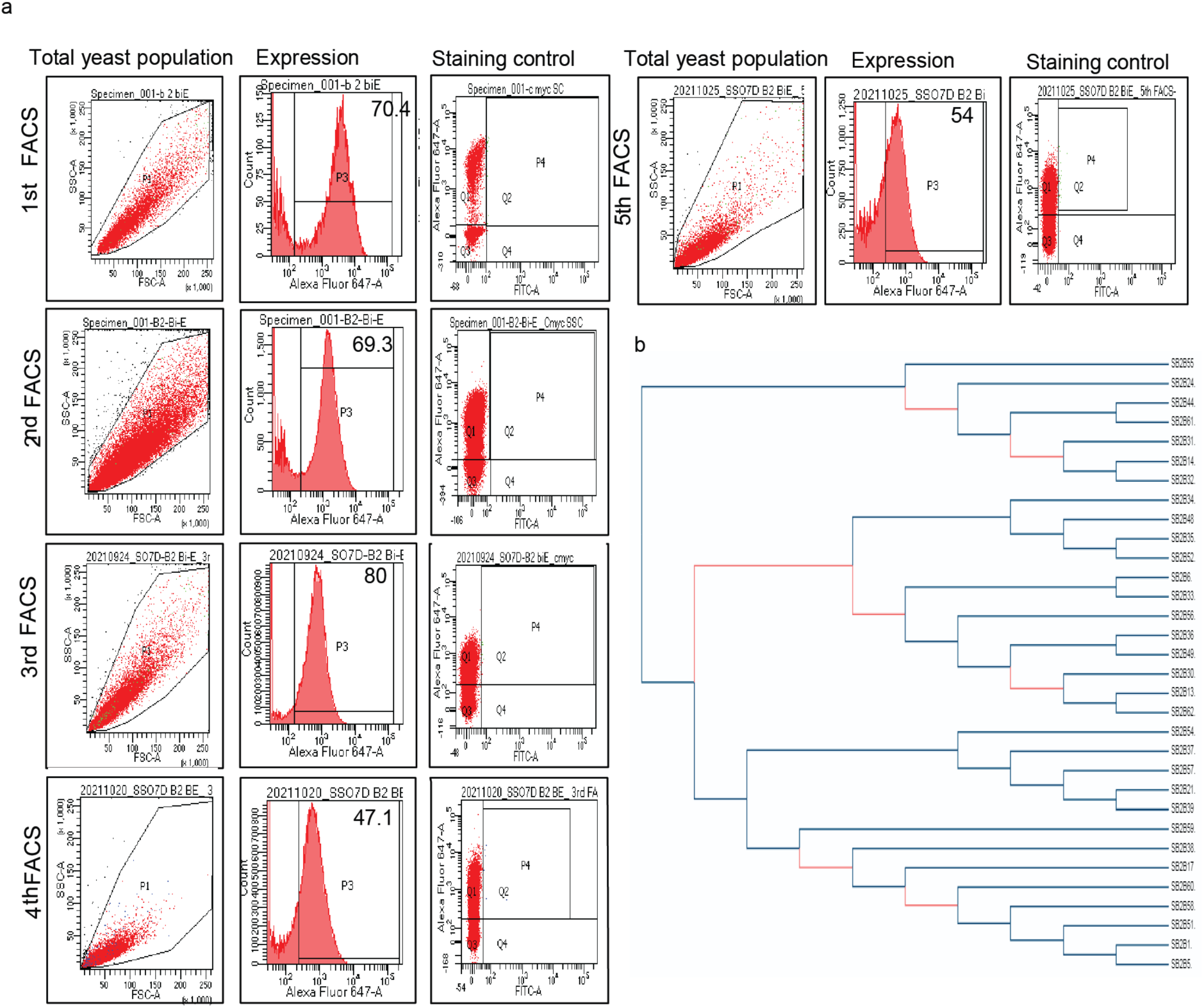
SSO7D Yeast display library screening and sequence enrichment. **a.** FACS plots showing the percentage of expressing clones for each FACS round, stained with Anti c-myc or HA, followed by anti-chicken or anti-rabbit Alexa Fluor 633, respectively (Methods). The gate for sorting the double-positive population in peptide-incubated library cultures was set using the third graph (staining control) in each FACS round. The library expression and peptide binding are displayed on the vertical and horizontal axes of the third graph, respectively. **b.** Phylogenetic tree of the individual SSO7D clones after the 5^th^ FACS round, illustrating the enrichment and diversity of potential glutamylation binder clones (mention total and unique clone numbers).

**Extended Data Fig 2.**
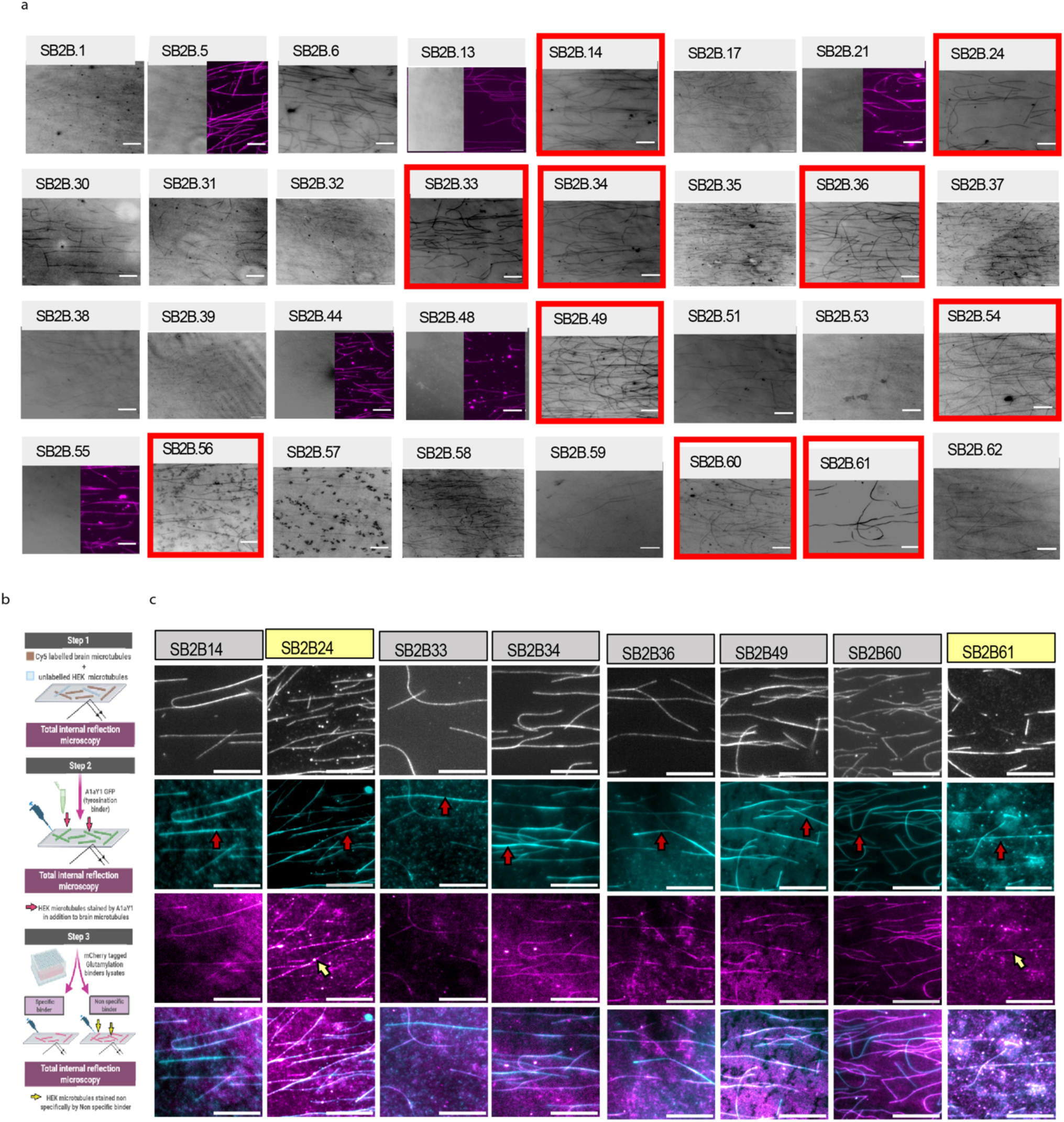
In vitro TIRFM-based assay for assessing the ability of SSO7D clones to bind to brain microtubules and HEK microtubules. **a.** TIRFM images of mCherry-SSO7D binders from E. coli lysates decorating goat brain microtubules. Only the mCherry-SSO7D binders that decorate microtubules are shown in grayscale. SSO7D clones that do not bind microtubules are displayed as a half field of view in grayscale, with Cy5-labeled microtubules in magenta. Red outlined boxes indicate strong microtubule SSO7D binders selected for further analysis. Scale bar: 10 μm. **b.** Schematic illustration of an in vitro TIRFM-based assay setup to assess the specificity of SSO7D clones toward brain and HEK microtubules. Step 1: Cy5-labeled goat microtubules and unlabeled HEK microtubules were mixed and flowed into the chamber. Step 2: A1aY1-GFP was added to label all microtubules. Step 3: Purified mCherry-SSO7D binders were introduced. SSO7D binders specific to glutamylated microtubules will label only goat brain microtubules, whereas non-specific binders will label both, illustrated as specific and non-specific binders. **c**. Cy5-labeled goat brain microtubules are shown in gray, while A1aY1-GFP-labeled microtubules are in cyan. HEK microtubules remain unlabeled but are visible through the A1aY1-GFP channel and are indicated by red arrows. Microtubules labeled by mCherry-SSO7D binders are shown in magenta. Non-specific binders (SB2B24 and SB2B61) that label HEK microtubules are marked with yellow arrows. Scale bar: 10 μm.

**Extended Data Fig. 3.**
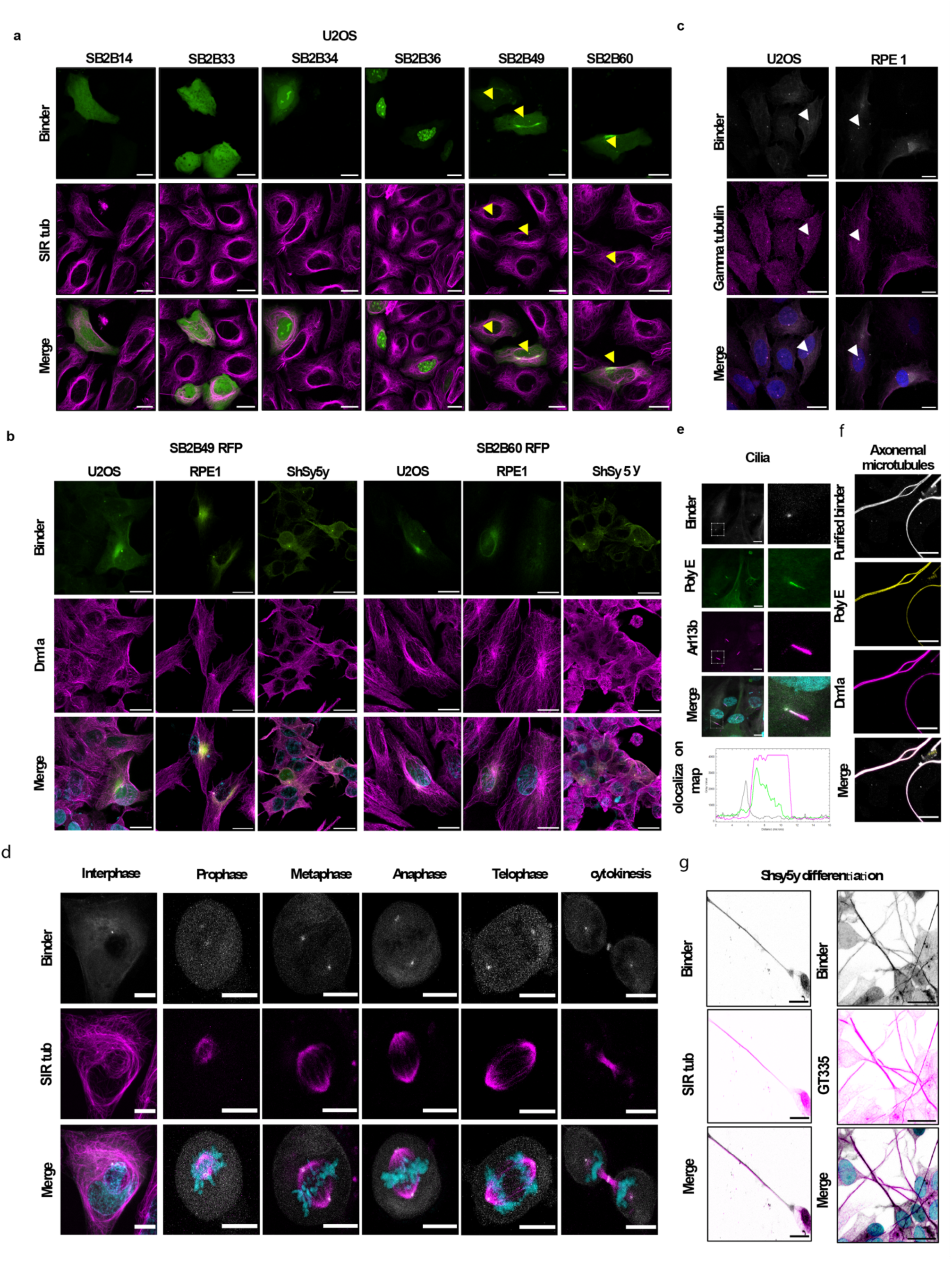
Binding properties of the glutamylation binder toward native glutamylation structures. **a.** Live cell confocal images of Tag RFP-T fused glutamylation binders transfected into U2OS cells. The cellular microtubules were visualized using a live cell microtubule marker, SIR tubulin (magenta), with the binders shown in green. The glutamylation binders (SB2B49 and SB2B60) that label the centrosomes are indicated by yellow arrows. Scale bar = 20 µm. **b.** Representative confocal images of U2OS, RPE1, and Shsy5y cell lines expressing SB2B49 and SB2B60. The cells were fixed and stained with the Dm1a antibody to visualize microtubules (magenta), and the binders are shown in green. Scale bar = 20 μm. **c.** Immunostaining of U2OS and RPE1 cells expressing Tag RFP-T fused to SB2B49 (grayscale) with anti-Gamma tubulin antibody (magenta) shows colocalization at the centrosomes. **d**. Live-cell imaging of microtubules and glutamylation across the cell cycle. Representative confocal images of different U2OS cells stably expressing SB2B49–TagRFP-T, captured in various cell-cycle stages as mentioned. Microtubules were visualized with SiR-tubulin (magenta); SB2B49 appears in grayscale; DNA was labeled with Hoechst (cyan). **e**. Ciliary microtubule glutamylation in RPE1 cells. Representative images of ciliated RPE1 cells stably expressing SB2B49–TagRFP-T (grayscale), fixed and stained for Arl13b (magenta) and PolyE (green). The line plot profile of cilia shows the extent of labeling for each marker. Scale bars = 10 µm. **f**. Axonemal microtubule glutamylation in mouse sperm. Isolated sperm were fixed, permeabilized, and stained with mCherry-tagged SB2B49 (green), alpha-tubulin (DM1A; magenta), and PolyE (yellow) to assess axoneme labeling. **g.** Axonal microtubule labeling during neuronal differentiation. Representative images of differentiated SH-SY5Y cells stably expressing SB2B49–TagRFP-T (grayscale). Microtubules were visualized with SiR-tubulin (magenta) in live cells and GT335 staining (magenta) in fixed cells, as indicated. Scale bars = 20 µm.

**Extended Data Fig 4.**
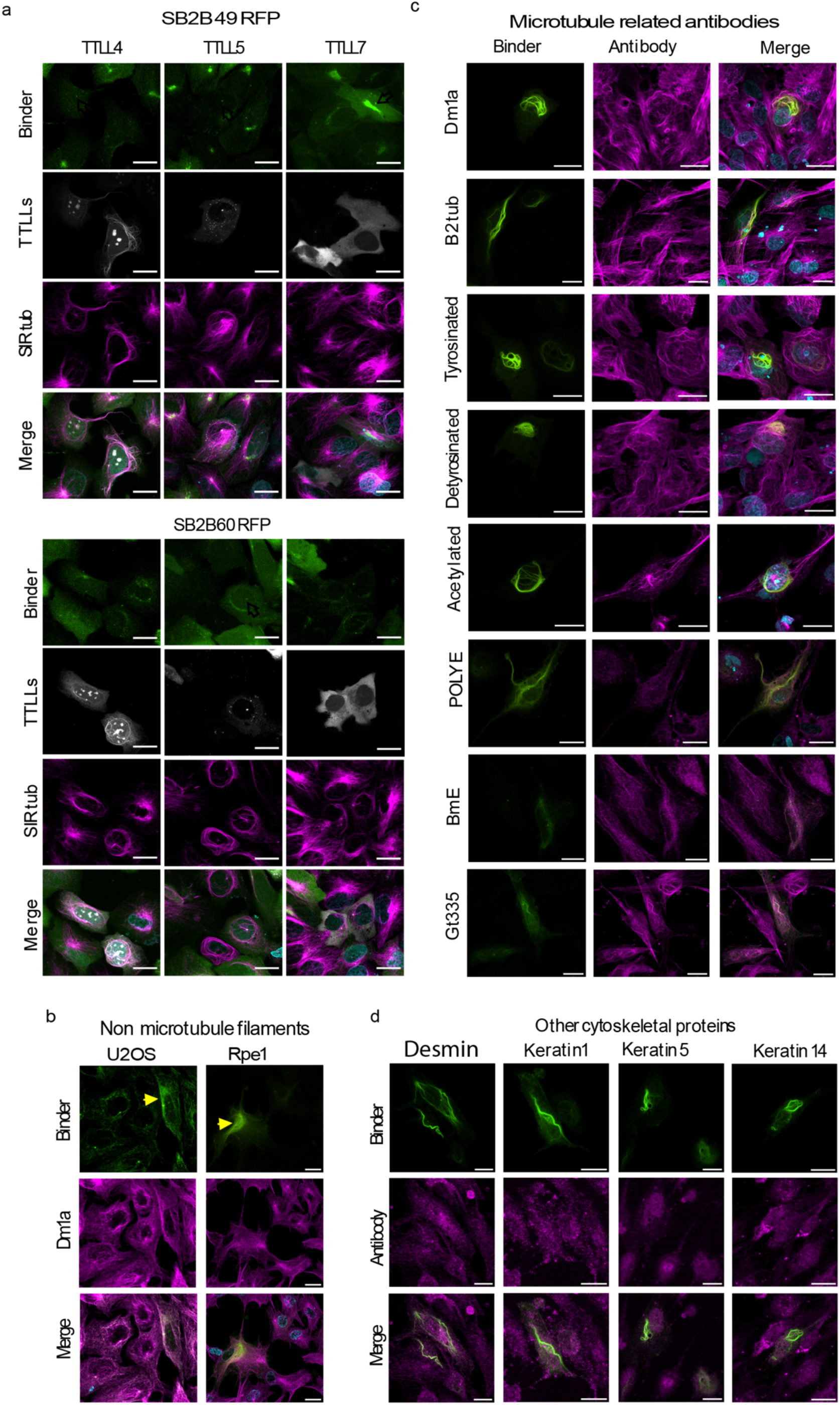
Glutamylation binds non-microtubule filamentous structures in cells. **a.** Representative live cell images of U2OS cells stably expressing two potential glutamylation sensors, SB2B49 and SB2B60, tagged with Tag RFP T (green), and transiently transfected with TTLL4, TTLL5, and TTLL7 fused to YFP (grey). Microtubules are visualized with SIR tubulin (magenta). Scale bar = 20 μm. **b**. Immunostaining of U2OS and RPE1 cells stably expressing Tag RFP-T fused to SB2B49 (green), with the DM1a antibody (magenta), shows binder labeling of non-microtubule filamentous structures. Scale bar = 20 μm. **C & d**. Confocal images of immunostained RPE1 cells stably expressing Tag RFP-T fused to SB2B49 (green), with various cytoskeleton and microtubule modification antibodies as indicated. Only the GT335 antibody staining overlaps with the SB2B49 binder fluorescence signal. Scale bar = 20 μm.

**Extended Data Fig 5.**
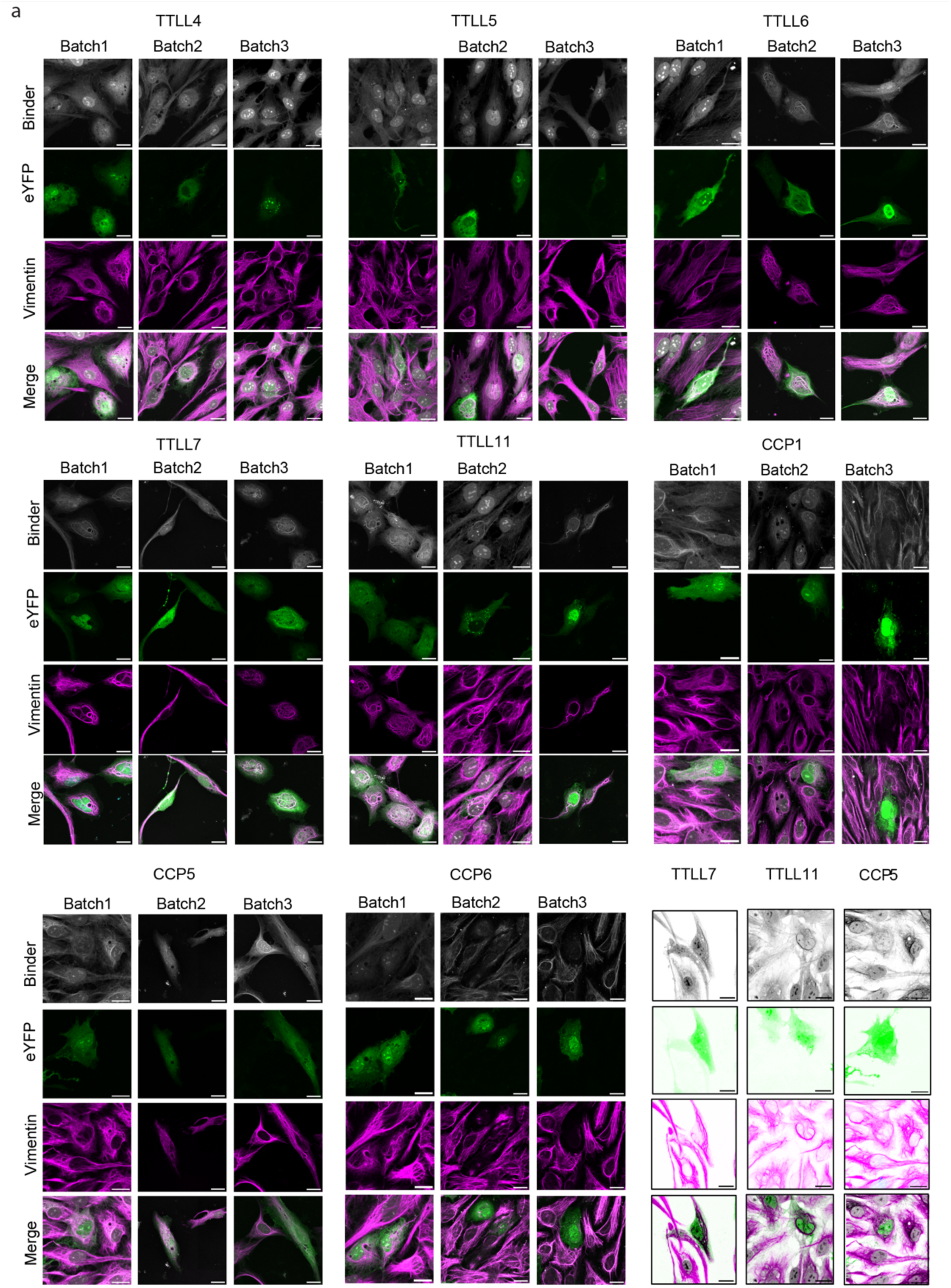
Effects of glutamylation-modifying enzymes on SB2B49 binding to vimentin filaments. **a.** Representative confocal images of immunostained RPE1 cells stably expressing Tag RFP-T fused to SB2B49 (magenta) and transiently transfected with TTLL4, TTLL5, TTLL6, TTLL7, TTLL11, CCP1, CCP5, and CCP6 e-YFP (green) against anti-vimentin antibodies (grayscale). Scale bar = 20 μm.

**Extended Data Fig 6.**
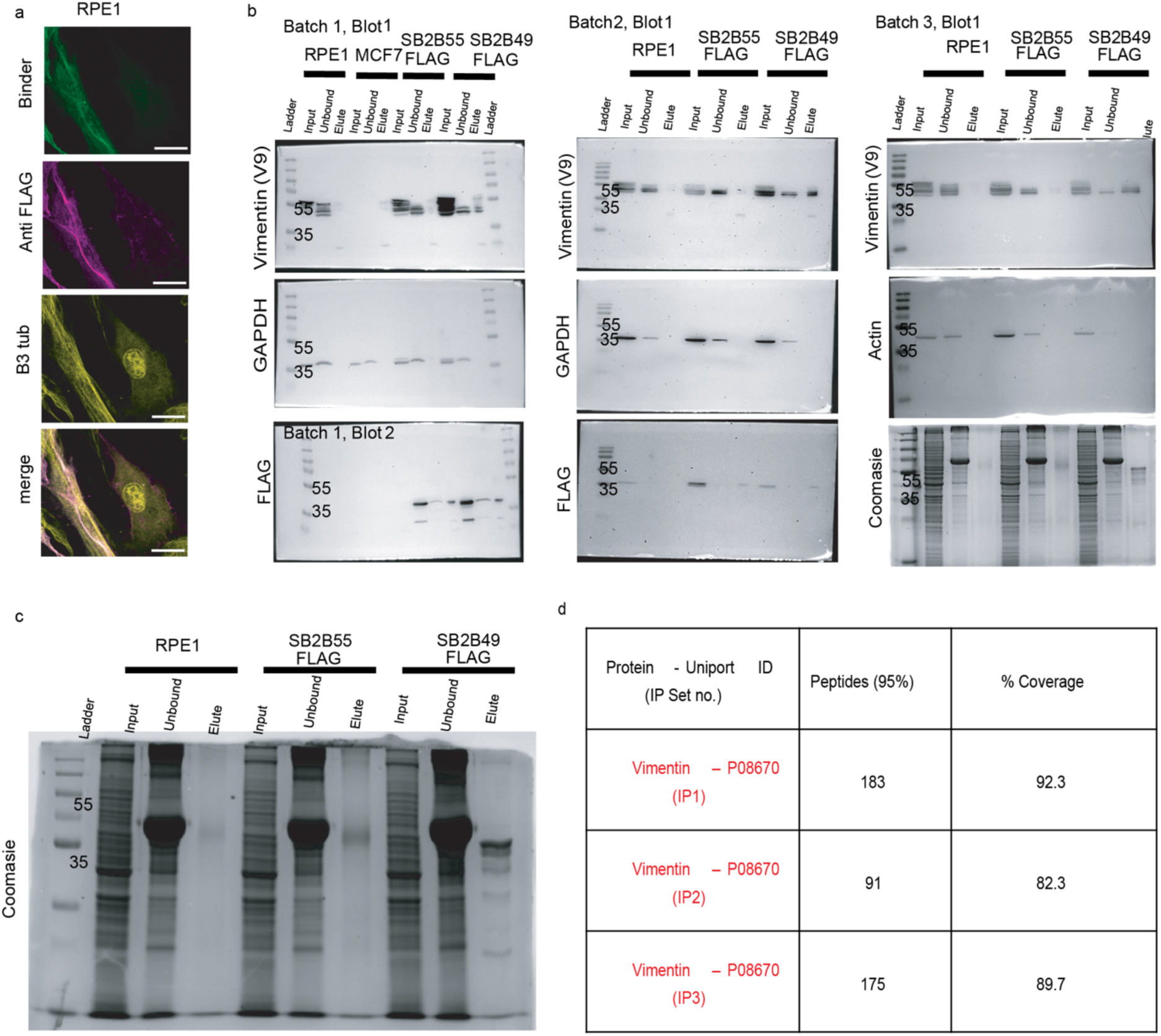
Biochemical validation of SB2B49 in recognizing glutamylated vimentin. **a.** Confocal images of immunostained RPE1 cells stably expressing Tag RFP-T fused to SB2B49-FLAG Tag (green) with anti-Flag (magenta) and anti-tubulin (yellow) antibodies as indicated. Scale bar = 20 μm. **b & c.** Uncropped Western blots and Coomassie-stained SDS-PAGE gel of pulldown with anti-Flag beads as indicated. A cropped image of Western blots from the second set and the Coomassie gel image shown in c are displayed in Figure 3. **d.** Summary of protein identification of pull-down samples using SB2B49 binder as shown in b & c (Methods).

**Extended Data Fig 7.**
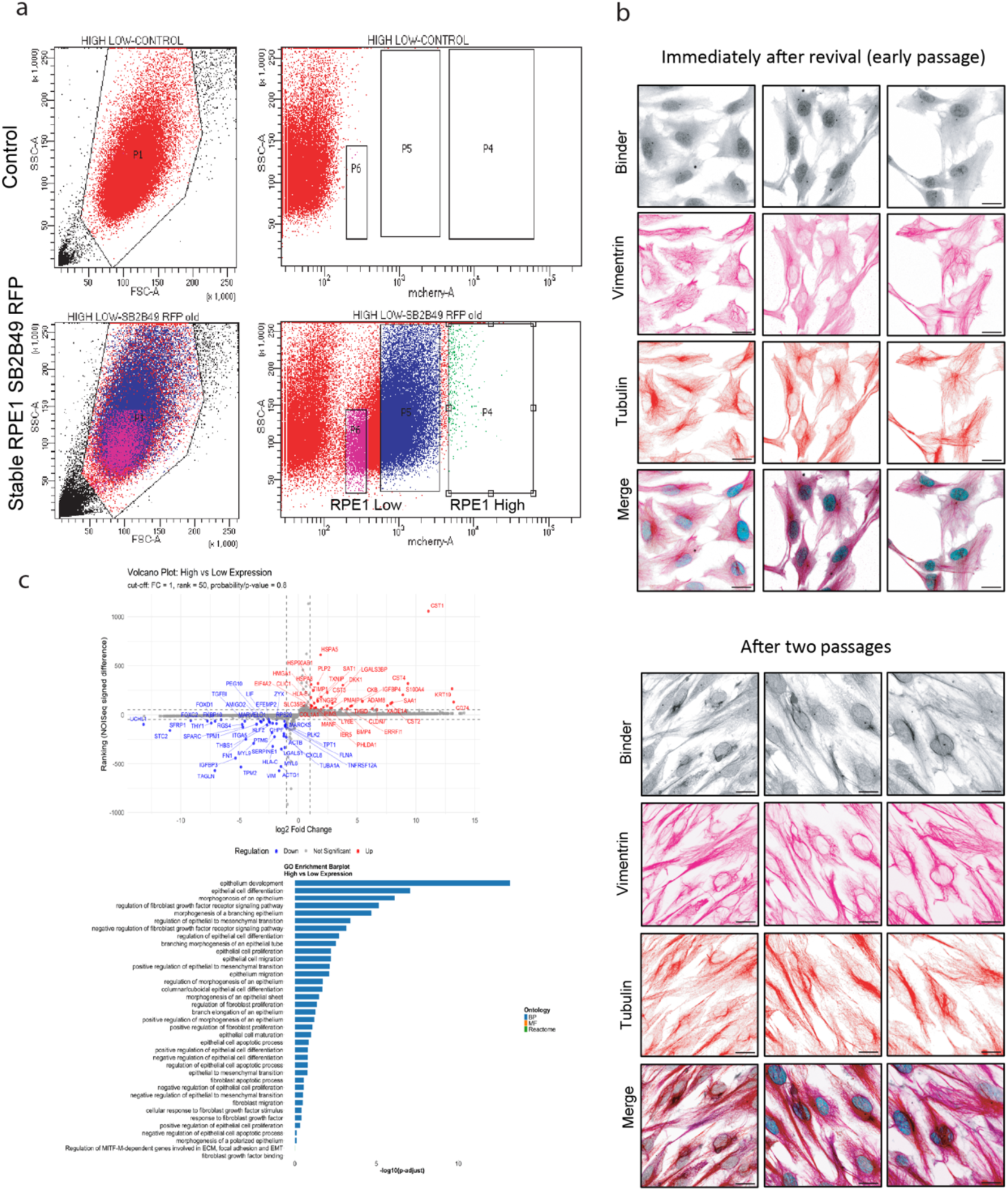
Glutamylated vimentin is linked to an epithelial-like cell state. **a.** RPE1 cells stably expressing N Tag RFP T fused SB2B49 binder were sorted into high and low expression *groups (bottom)* using wild-type RPE1 as a control *(top).* The first plot for both the control and stable RPE1 cells displays the total cell population, which was gated for sorting. The stable RPE1 cells were sorted into high (RPE1 High) and low (RPE1 Low) expression based on the gating in the second control FACS plot. **b**. Wild-type RPE1 cells from three different batches after immediate revival (RPE_fresh-seed_) and the respective batches after two passages (RPE_passaged_), stained with recombinant SB2B49-CF640R (grayscale), vimentin (magenta), and DM1a (red) antibodies, show the emergence of a glutamylated vimentin form after two passages. Scale bar = 20 µm. **c**. Gene Ontology and volcano plot of transcriptomic data for RPE_fresh-seed_ versus RPE_passaged_ show an epithelial-like state in RPE_passaged_ cells.

**Extended Data Fig 8.**
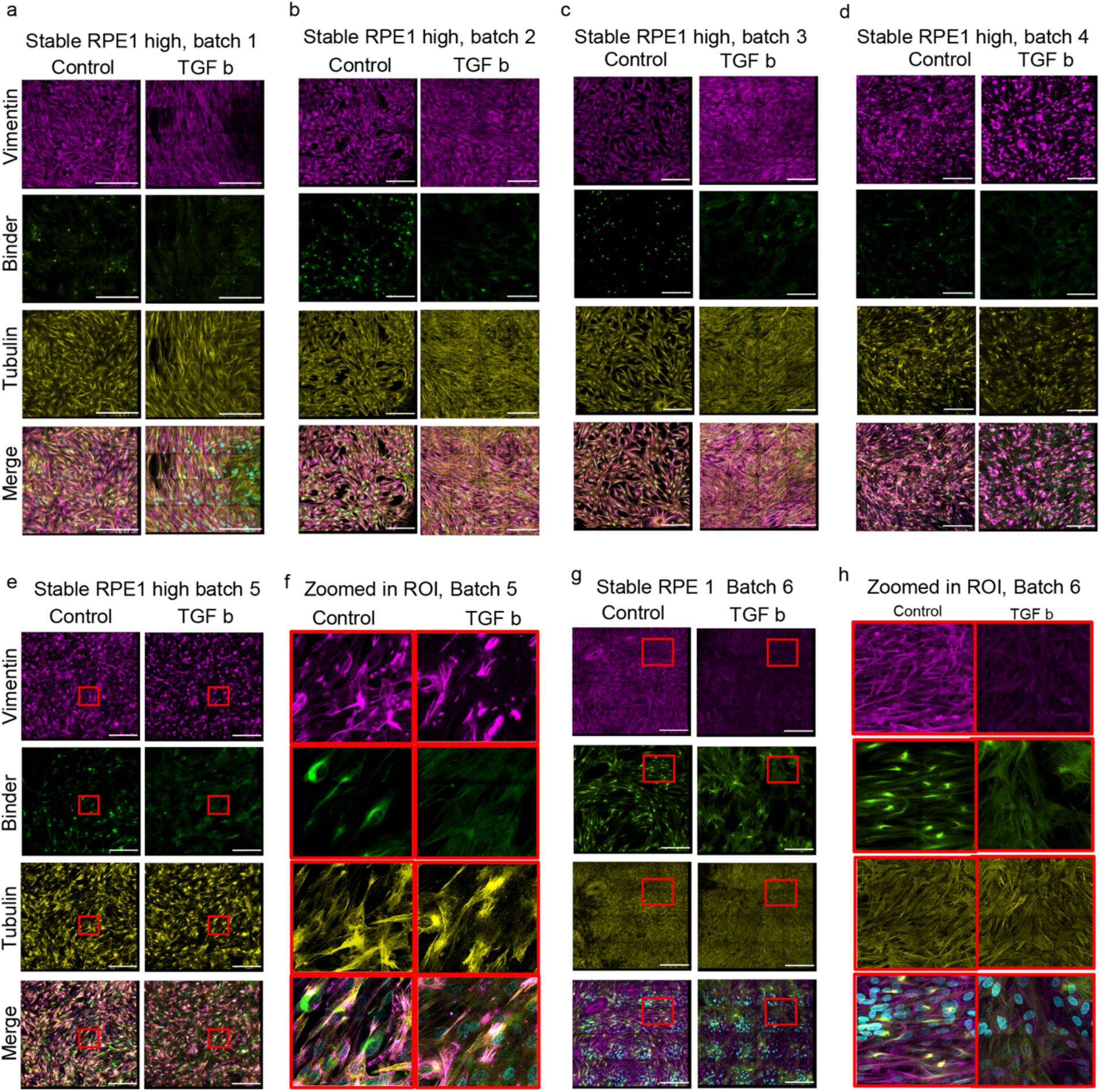
Regulation of glutamylated vimentin with respect to external stimuli. **a-h.** Confocal tiled images (4x4 FOVs) of experiments from batches 1-4 showing RPE1 cells stably expressing N Tag RFP T fused SB2B49 binder (green), compared between untreated (Control) and TGF-beta treated samples, stained with vimentin (magenta) and DM1a (yellow). Only images from batches 5 and 6 are highlighted with a zoomed ROI (red dotted boxes). Scale bars: a-d, e, and g = 200 µm.

**Extended Data Fig 9.**
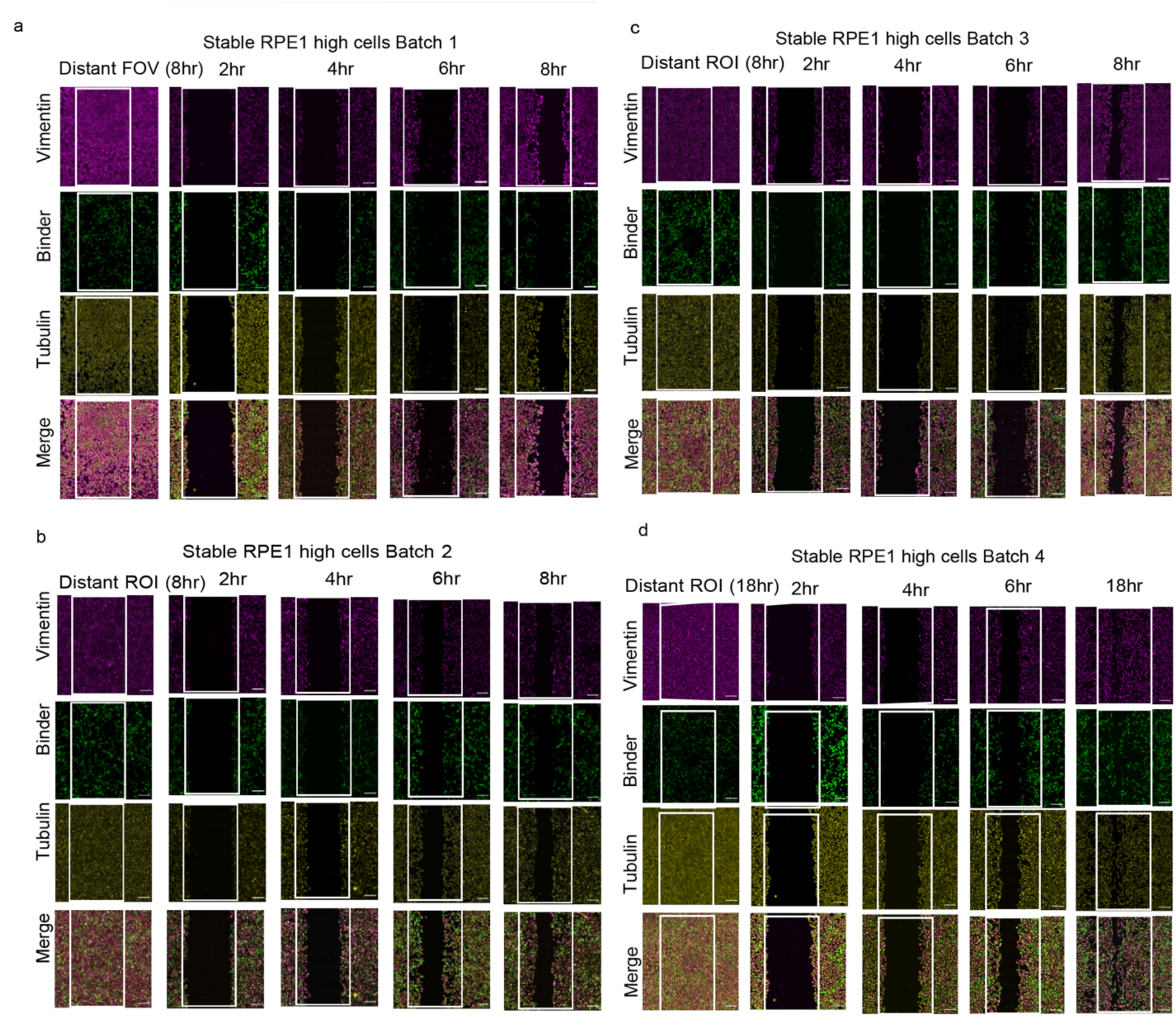
Effects of glutamylated vimentin during cell migration. **a-c.** Confocal tiled images from batches 1-3 used in the quantification graph shown in Figure 6, featuring RPE1 cells stably expressing N-Tag RFP-T fused SB2B49 (green), stained with vimentin (magenta) and DM1a (yellow). The white dotted box indicates the scratch area created with a pipette tip. Two 8x8 tiled images were captured for each dish: one from the scratch zone and one from a distant region (DISTANT ROI). Only the 8-hour DISTANT ROI image is shown for reference. The second and third quantification batches are displayed in panels **b** and **c,** respectively. **d.** Confocal tiled images from batch 4. Scale bar = 200 μm.

**Extended Data Fig 10.**
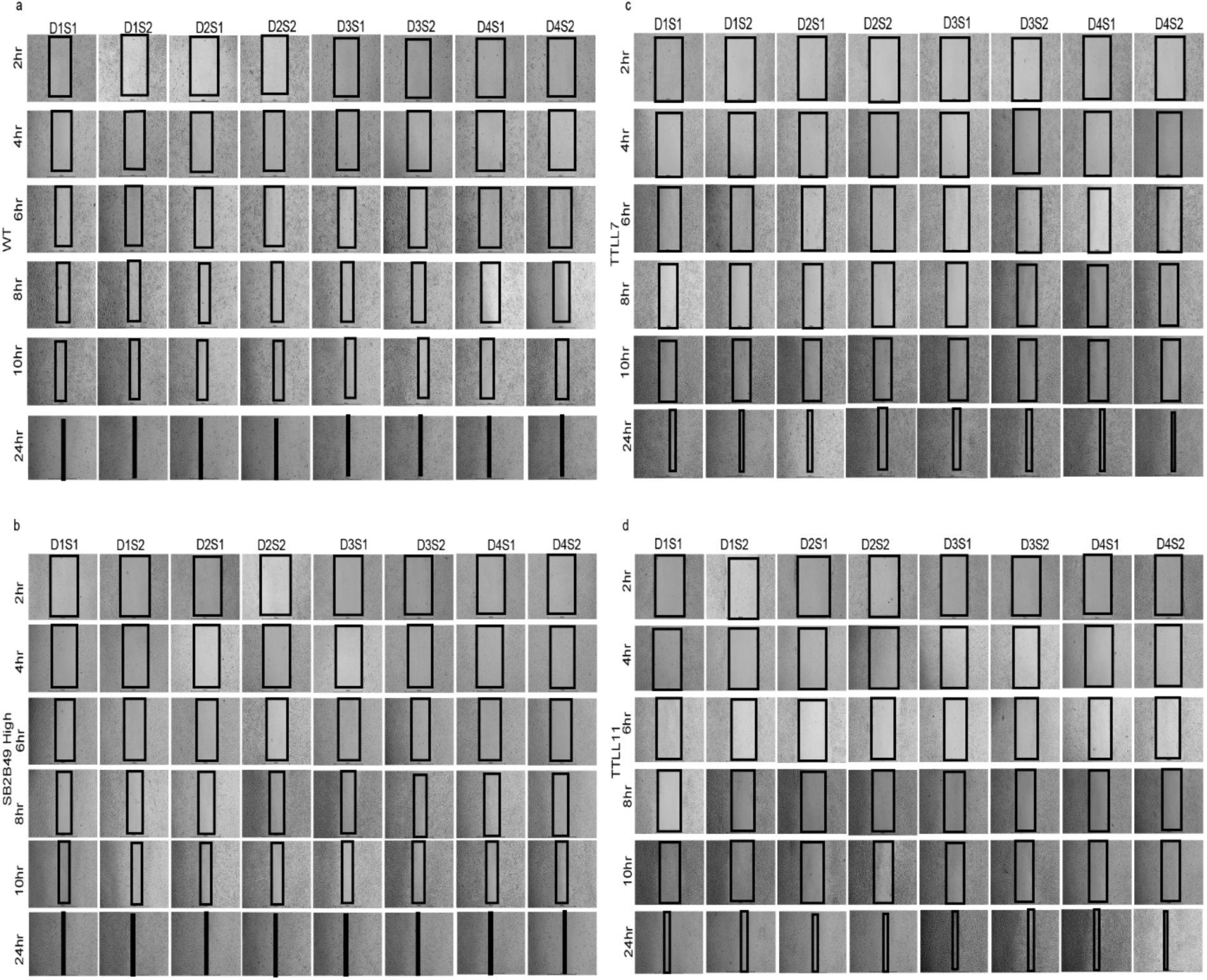
Cells with glutamylated vimentin show reduced migration capacity. **a-d**. Bright-field images of RPE1 wild-type cells and cells stably expressing either N Tag RFP T fused SB2B49 binder or TTLL7-eYFP or TTLL11-eYFP from four different culture dishes, with two scratches made in each dish; ‘D’ and ‘S’ represent the dish number and scratch number from the respective dish. The scratches were generated as described in Extended Data Fig 9 and Methods. Scale bar = 1000 µm.

## References

1. Fletcher, D. A. & Mullins, R. D. Cell mechanics and the cytoskeleton. Nature 463, 485–492 (2010).

2. Choi, S. R. et al. Structural basis of microtubule-mediated signal transduction. Cell 189, 461–477.e16 (2026).

3. Janke, C. & Magiera, M. M. The tubulin code and its role in controlling microtubule properties and functions. Nat. Rev. Mol. Cell Biol. 21, 307–326 (2020).

4. L’Hernault, S. W. & Rosenbaum, J. L. Chlamydomonas alpha-tubulin is posttranslationally modified by acetylation on the epsilon-amino group of a lysine. Biochemistry 24, 473–478 (1985).

5. Barra, H. S., Rodriguez, J. A., Arce, C. A. & Caputto, R. A soluble preparation from rat brain that incorporates into its own proteins ( 14 C)arginine by a ribonuclease-sensitive system and ( 14 C)tyrosine by a ribonuclease-insensitive system. J. Neurochem. 20, 97–108 (1973).

6. Aillaud, C. et al. Vasohibins/SVBP are tubulin carboxypeptidases (TCPs) that regulate neuron differentiation. Science 358, 1448–1453 (2017).

7. Eddé, B. et al. Posttranslational glutamylation of alpha-tubulin. Science 247, 83–85 (1990).

8. Sirajuddin, M., Rice, L. M. & Vale, R. D. Regulation of microtubule motors by tubulin isotypes and post-translational modifications. Nat. Cell Biol. 16, 335–344 (2014).

9. McKenney, R. J., Huynh, W., Vale, R. D. & Sirajuddin, M. Tyrosination of α-tubulin controls the initiation of processive dynein-dynactin motility. EMBO J. 35, 1175–1185 (2016).

10. Genova, M. et al. Tubulin polyglutamylation differentially regulates microtubule-interacting proteins. EMBO J. 42, e112101 (2023).

11. Barisic, M. et al. Mitosis. Microtubule detyrosination guides chromosomes during mitosis. Science 348, 799–803 (2015).

12. Magiera, M. M. et al. Excessive tubulin polyglutamylation causes neurodegeneration and perturbs neuronal transport. EMBO J. 37, e100440 (2018).

13. Terman, J. R. & Kashina, A. Post-translational modification and regulation of actin. Curr. Opin. Cell Biol. 25, 30–38 (2013).

14. Varland, S., Vandekerckhove, J. & Drazic, A. Actin Post-translational Modifications: The Cinderella of Cytoskeletal Control. Trends Biochem. Sci. 44, 502–516 (2019).

15. Herrmann, H. & Aebi, U. Intermediate Filaments: Structure and Assembly. Cold Spring Harb. Perspect. Biol. 8, a018242 (2016).

16. Etienne-Manneville, S. Cytoplasmic Intermediate Filaments in Cell Biology. Annu. Rev. Cell Dev. Biol. 34, 1–28 (2018).

17. Coelho-Rato, L. S., Parvanian, S., Andrs Salajkova, S., Medalia, O. & Eriksson, J. E. Intermediate filaments at a glance. J. Cell Sci. 137, jcs261386 (2024).

18. Kalluri, R. & Weinberg, R. A. The basics of epithelial-mesenchymal transition. J. Clin. Invest. 119, 1420–1428 (2009).

19. Lamouille, S., Xu, J. & Derynck, R. Molecular mechanisms of epithelial-mesenchymal transition. Nat. Rev. Mol. Cell Biol. 15, 178–196 (2014).

20. Guo, M. et al. Vimentin intermediate filaments as structural and mechanical coordinators of mesenchymal cells. Nat. Cell Biol. 27, 1210–1218 (2025).

21. Thiery, J. P., Acloque, H., Huang, R. Y. J. & Nieto, M. A. Epithelial-mesenchymal transitions in development and disease. Cell 139, 871–890 (2009).

22. Pastushenko, I. & Blanpain, C. EMT Transition States during Tumor Progression and Metastasis. Trends Cell Biol. 29, 212–226 (2019).

23. Jolly, M. K., Ware, K. E., Gilja, S., Somarelli, J. A. & Levine, H. EMT and MET: necessary or permissive for metastasis? Mol. Oncol. 11, 755–769 (2017).

24. Nieto, M. A., Huang, R. Y.-J., Jackson, R. A. & Thiery, J. P. EMT: 2016. Cell 166, 21–45 (2016).

25. Yang, J. et al. Guidelines and definitions for research on epithelial–mesenchymal transition. Nat. Rev. Mol. Cell Biol. 21, 341–352 (2020).

26. Hyder, C. L., Isoniemi, K. O., Torvaldson, E. S. & Eriksson, J. E. Insights into intermediate filament regulation from development to ageing. J. Cell Sci. 124, 1363–1372 (2011).

27. Janke, C. et al. Tubulin polyglutamylase enzymes are members of the TTL domain protein family. Science 308, 1758–1762 (2005).

28. Kalinina, E. et al. A novel subfamily of mouse cytosolic carboxypeptidases. FASEB J. 21, 836–850 (2007).

29. van Dijk, J. et al. Polyglutamylation is a post-translational modification with a broad range of substrates. J. Biol. Chem. 283, 3915–3922 (2008).

30. Rogowski, K. et al. A family of protein-deglutamylating enzymes associated with neurodegeneration. Cell 143, 564–578 (2010).

31. Wolff, A. et al. Distribution of glutamylated alpha and beta-tubulin in mouse tissues using a specific monoclonal antibody, GT335. Eur. J. Cell Biol. **59**, 425–432 (1992).

32. Bodakuntla, S., Janke, C. & Magiera, M. M. Tubulin polyglutamylation, a regulator of microtubule functions, can cause neurodegeneration. Neurosci. Lett. 746, 135656 (2021).

33. van Dijk, J. et al. A targeted multienzyme mechanism for selective microtubule polyglutamylation. Mol. Cell 26, 437–448 (2007).

34. Kesarwani, S. et al. Genetically encoded live-cell sensor for tyrosinated microtubules. J. Cell Biol. 219, e201912107 (2020).

35. Rüdiger, M., Plessman, U., Klöppel, K. D., Wehland, J. & Weber, K. Class II tubulin, the major brain beta tubulin isotype is polyglutamylated on glutamic acid residue 435. FEBS Lett. 308, 101–105 (1992).

36. Souphron, J. et al. Purification of tubulin with controlled post-translational modifications by polymerization-depolymerization cycles. Nat. Protoc. 14, 1634–1660 (2019).

37. Janke, C. The tubulin code: molecular components, readout mechanisms, and functions. J. Cell Biol. 206, 461–472 (2014).

38. Thazhath, R., Liu, C. & Gaertig, J. Polyglycylation domain of beta-tubulin maintains axonemal architecture and affects cytokinesis in Tetrahymena. Nat. Cell Biol. 4, 256–259 (2002).

39. Pieper, F. R. et al. Regulation of vimentin expression in cultured epithelial cells. Eur. J. Biochem. 210, 509–519 (1992).

40. Usman, S. et al. Transcriptome Analysis Reveals Vimentin-Induced Disruption of Cell–Cell Associations Augments Breast Cancer Cell Migration. Cells 11, 4035 (2022).

41. Bhowmick, N. A. et al. Transforming Growth Factor-β1 Mediates Epithelial to Mesenchymal Transdifferentiation through a RhoA-dependent Mechanism. Mol. Biol. Cell 12, 27–36 (2001).

42. Mendez, M. G., Kojima, S.-I. & Goldman, R. D. Vimentin induces changes in cell shape, motility, and adhesion during the epithelial to mesenchymal transition. FASEB J. 24, 1838–1851 (2010).

43. Usman, S. et al. Vimentin Is at the Heart of Epithelial Mesenchymal Transition (EMT) Mediated Metastasis. Cancers 13, 4985 (2021).

44. Yu, I., Garnham, C. P. & Roll-Mecak, A. Writing and Reading the Tubulin Code. J. Biol. Chem. 290, 17163–17172 (2015).

45. Eriksson, J. E. et al. Specific in vivo phosphorylation sites determine the assembly dynamics of vimentin intermediate filaments. J. Cell Sci. 117, 919–932 (2004).

46. Chou, Y. H., Rosevear, E. & Goldman, R. D. Phosphorylation and disassembly of intermediate filaments in mitotic cells. Proc. Natl. Acad. Sci. U. S. A. 86, 1885–1889 (1989).

47. Gera, N., Hill, A. B., White, D. P., Carbonell, R. G. & Rao, B. M. Design of pH sensitive binding proteins from the hyperthermophilic Sso7d scaffold. PloS One 7, e48928 (2012).

